# Genome-wide distribution of Rad26 and Rad1-Rad10 reveals their relationship with Mediator and RNA polymerase II

**DOI:** 10.1101/2021.10.20.465181

**Authors:** Diyavarshini Gopaul, Cyril Denby Wilkes, Arach Goldar, Nathalie Giordanengo Aiach, Marie-Bénédicte Barrault, Elizaveta Novikova, Julie Soutourina

## Abstract

Transcription is coupled with DNA repair, especially within nucleotide excision repair (NER). Mediator is a conserved coregulator playing a key role in RNA polymerase (Pol) II transcription. Mediator also links transcription and NER via a direct contact with Rad2/XPG endonuclease. In this work, we analyzed the genome-wide distribution of Rad26/CSB and that of Rad1-Rad10/XPF-ERCC1, addressing the question on a potential interplay of these proteins with Mediator and Pol II in yeast *Saccharomyces cerevisiae*. Our genome-wide analyses show that Rad1-Rad10 and Rad26 are present on the yeast genome in the absence of genotoxic stress, especially on highly transcribed regions, with Rad26 binding strongly correlating with that of Pol II. Moreover, we revealed that Rad1-Rad10 and Rad26 colocalize with Mediator on intergenic regions and physically interact with this complex. Using *kin28* TFIIH mutant, we showed that Mediator stabilization on core promoters lead to an increase in Rad1-Rad10 chromatin binding, whereas Rad26 occupancy is less impacted by Mediator and follows mainly a decrease in Pol II transcription. Combined with multivariate analyses, our results reveal the interplay between Rad1-Rad10, Rad26, Mediator and Pol II, modulated by the binding dynamics of Mediator and Pol II transcription. In conclusion, we extend the Mediator link to Rad1-Rad10 and Rad26 NER proteins and reveal important differences in Mediator relationships with Rad2, Rad1-Rad10 and Rad26. Our work thus contributes to new concepts of the functional interplay between transcription and DNA repair, relevant for human diseases including cancer and XP/CS syndromes.

## Introduction

Chromatin binding constitutes an essential step for the functions of nuclear proteins. Genome-wide location analysis is thus a powerful tool to understand *in vivo* mechanisms of the chromatin-associated proteins involved in transcription and DNA repair processes. These analyses revealed the chromatin binding of Rad2, nucleotide excision DNA repair (NER) protein, in the absence of exogenous genotoxic stress (Eyboulet et al. 2013). A high correlation between Rad2 and Mediator occupancies of regulatory regions (upstream activating sequences, UAS) was one of the starting points to propose functional interplay between these two nuclear components. NER constitutes of two mechanistically distinct subpathways: global genome repair (GGR) that occurs on chromatin with different functional states and compaction levels, along with transcription-coupled repair (TCR) that corresponds to a specialized pathway allowing efficient repair of Pol II-blocking DNA damages and resumption of transcription (Fousteri and Mullenders 2008; Hanawalt and Spivak 2008). The NER repair machinery consists of near 30 proteins, but genome-wide distribution of many NER proteins remains unknown in the absence of exogeneous genotoxic stress, as well as their relationship with Mediator and Pol II.

Rad26 in yeast *S. cerevisiae*, homologous to Cockayne Syndrome CSB protein in mammalian cells, plays an essential role in the TCR pathway by interacting with Pol II to initiate the TCR complex assembly (reviewed in (Mullenders 2015; Boetefuer et al. 2018)). Several lines of evidence pointed out that Rad26/CSB activity is closely related to Pol II transcription. Cryo-EM structures of yeast Rad26 – Pol II (Xu et al. 2017) and human Pol II complexes with CSB and other TCR factors (Kokic et al. 2021) were recently determined, providing mechanistic insights into Rad26 role in TCR and in the resolution of Pol II stalling in general (Wang et al. 2018). Transcription-dependent binding of Rad26 was observed on selected transcribed regions for two inducible genes and one constitutively-expressed gene without exogeneous genotoxic stress (Malik et al. 2010). In human cells, ChIP-seq analysis also showed CSB association to chromatin in the absence of UV irradiation (Lake et al. 2014; Wang et al. 2014). Fluorescence imaging experiments showed a dynamic association of CSB with chromatin before UV irradiation and a stabilization of this association upon UV stress (van den Boom et al. 2004). Several lines of evidence were provided for CSB implication in transcription after UV irradiation (Rockx et al. 2000; Proietti-De-Santis et al. 2006; Epanchintsev et al. 2017) or during replicative cell growth (Lake et al. 2014; Wang et al. 2014). *In vitro*, CSB was shown to enhance Pol II transcription elongation (Selby and Sancar 1997a). CSB interacts with stalled Pol II complex and promotes transcription at natural pause sites (Sarker et al. 2005).

Rad1-Rad10 in yeast *S. cerevisiae*, homologous to XPF-ERCC1 mammalian complex, is a structure-specific endonuclease essential for NER pathway (reviewed in (McNeil and Melton 2012; Faridounnia et al. 2018)). Rad1-Rad10/XPF-ERCC1 functions as endonuclease only as a heterodimer in which Rad1/XPF provides the 5’-endonuclease activity and Rad10/ERCC1 regulates DNA- and protein-protein interactions. *In vitro*, the two proteins are unstable in the absence of each other (Bailly et al. 1992; van Vuuren et al. 1993). Together with Rad2/XPG 3’-endonuclease, Rad1-Rad10/XPF-ERCC1 is essential for the dual incision of the damaged DNA in both NER pathways. Mutations in XPF-ERCC1 human genes led to xeroderma pigmentosum (XP) or XP combined with Cockayne syndrome (XP/CS), more severe XPF-ERCC1 progeroid syndrome (XF-E) or Fanconi Anemia (FA) (Cleaver et al. 2009; Menck and Munford 2014; Bukowska and Karwowski 2018). It has been recently shown that impaired NER and persistence of the repair machinery at DNA lesions characterize XPF XP/CS patients with additional developmental and neurodegenerative symptoms (Sabatella et al. 2018). A recent study reported that a dietary restriction in ERCC1-deficient mice with accelerated ageing features delayed genomic stress and improved their lifespan, thereby proposing a model for age-related diseases in which DNA lesions from exogenous and endogenous sources reduce transcriptional output in a gene-size-dependent manner (Vermeij et al. 2016). Additional roles of XPF-ERCC1 were also proposed in chromatin looping and optimal transcription activation of nuclear-receptor (NR)-dependent genes together with XPG and other NER factors (Le May et al. 2010; Le May et al. 2012). In mouse model, XPF-ERCC1 was dispensable for ongoing transcription, but was proposed to be involved in fine-turning of optimal activation of hepatic genes (Kamileri et al. 2012).

Mediator of transcriptional regulation is an essential multi-subunit coregulator playing a key role in RNA polymerase (Pol) II transcription (reviewed in (Kornberg 2005; Soutourina 2018)). This complex, conserved from yeast to human cells, is crucial to transmit regulatory information from specific transcription factors to Pol II basal transcription machinery. Importance of Mediator complex is highlighted by the fact that mutations or changes in the expression level of Mediator subunits was implicated in many human diseases including cancers or neurodevelopmental diseases (Spaeth et al. 2011; Schiano et al. 2014). Our previous work suggests that Mediator function extends beyond the transcription process (Eyboulet et al. 2013). We demonstrated that the Mediator complex links transcription and nucleotide excision DNA repair (NER) via Rad2/XPG 3’-endonuclease (Eyboulet et al. 2013). Mediator physically interacts with Rad2/XPG DNA repair protein (Eyboulet et al. 2013; Kikuchi et al. 2015). Genome-wide location analyses revealed Rad2 chromatin binding in the absence of exogenous genotoxic stress. Rad2 was associated with upstream activating sequences (UAS) and transcribed regions of Pol II-transcribed genes, as well as with Pol III-transcribed genes and telomeric regions (Eyboulet et al. 2013). No major transcriptional role was observed for Rad2 in yeast, but our findings suggested Mediator involvement in TCR. Genetic and genomic approaches further contributed to provide insights into the functional interplay between Rad2, Mediator and Pol II, suggesting that dynamic interactions between these nuclear components are involved in Rad2 loading to the chromatin (Georges et al. 2019).

In this study, we analyzed the genome-wide location of TCR-specific Rad26 protein and Rad1/Rad10 endonuclease that works together with Rad2 in NER. We addressed the question on the potential interplay of these proteins with Mediator and Pol II in the absence of genotoxic stress.

## Results

### Genome-wide location analysis of Rad26

To date, no genomic data are available for many NER proteins including Rad26, the TCR-specific component. To determine how this protein is distributed on the yeast chromatin, we performed Rad26 ChIP-seq experiments and analysis. Two yeast strains carrying N-terminal or C-terminal HA-tagged version of Rad26 were constructed. An increase in UV sensitivity was observed for the strain carrying C-terminally tagged Rad26 in a GGR-deficient *rad7Δ* context (**Supplementary Figure S1A**), suggesting that this version was not fully functional. The UV sensitivity was needed to be tested in a GGR-deficient context, since *rad26* deletion alone does not lead to UV sensitivity in yeast (van Gool et al. 1994). The N-terminal tagged version was therefore used for further study.

ChIP followed by qPCR on selected regions showed that Rad26 was enriched inside Pol II-transcribed genes (**Supplementary Figure S2A**). Specific examples of ChIP-seq profiles on **Figure 1A** illustrate Rad26 enrichment mostly on Pol II-transcribed genes and its binding on intergenic regions. Analysis of Rad26 enrichment peaks revealed that, surprisingly, this protein is also present on other genomic regions, in particular it is highly enriched on all 16 yeast centromeres (**Supplementary Figure S3A, B**). We compared genome-wide occupancy of Rad26 with that of Pol II on Pol II-transcribed regions and observed a linear relationship with a high R^2^ = 0.94 (**Figure 1B**) and Spearman correlation of 0.91. Heatmaps of tag density were generated for Rad26, Mediator (Med17 subunit) and Pol II and compared on Pol II-transcribed regions, clearly illustrating Rad26 colocalization with Pol II (**Figure 1C**).

**Figure 1.**
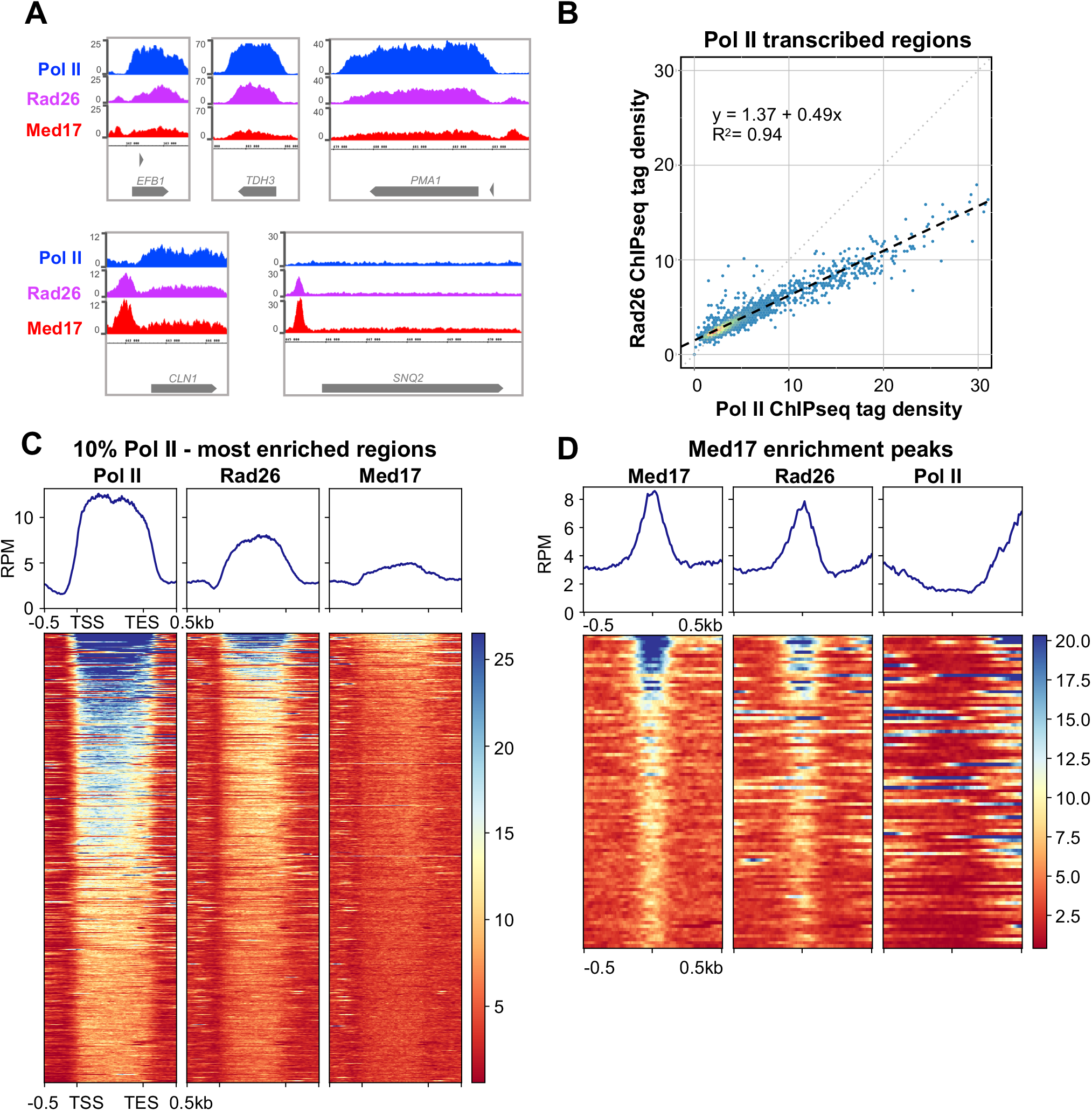
Genome-wide analysis of Rad26 occupancy. **(A)** Examples of Rad26, Pol II and Mediator (Med17) tag density profiles in WT context on Pol II-transcribed genes. **(B)** Rad26 ChIP-seq density versus Pol II ChIP-seq density on Pol II transcribed genes. Each point on the plot corresponds to one transcribed region. A linear regression (dotted line) and an R^2^ linear regression coefficient are indicated. The dashed line corresponds to y = x. **(C)** Heatmaps of Pol II, Rad26 and Mediator (Med17) ChIP-seq occupancy on 10% Pol II-most enriched regions (scaled windows for 500 bp before TSS, between TSS and TES, and 500 bp after TES), sorted by decreasing Pol II occupancy. Median tag density profiles in RPM (reads per million) are shown in upper panels. **(D)** Heatmaps of Mediator (Med17), Rad26 and Pol II ChIP-seq occupancy centered on Mediator enrichment peaks (−500 bp to +500 bp), sorted by decreasing Mediator occupancy. Median tag density profiles in RPM (reads per million) are shown in upper panels.

Further analysis revealed the presence of Rad26 on intergenic regions enriched by the Mediator complex. Spearman correlation analysis on the Mediator enrichment peaks showed a correlation between the two sets of data (Spearman correlation coefficient equal to 0.67). Heatmaps of tag density for Rad26, Mediator (Med17 subunit) and Pol II centered on Mediator enrichment peaks, as well as metagene analysis illustrated colocalization of Rad26 and Mediator (**Figure 1D**).

Previously, Rad26 occupancy of selected Pol II-transcribed genes without exogeneous genotoxic stress was shown to be transcription-dependent (Malik et al. 2010). Rad26 was enriched on transcribed regions of *GAL* and *INO1* genes under inducible conditions and *RPS5* constitutively-expressed gene (Malik et al. 2010). In addition, Rad26 association to *GAL1* gene upon galactose induction was reduced in *rpb1-1* Pol II mutant (Malik et al. 2010). A *rad26* deletion reduced Pol II occupancy only during induction of *GAL* genes, but did not affect the steady state level of the enzyme association upon full induction of *GAL* genes or constitutively-expressed *RPS5* gene (Malik et al. 2012). This suggests that Rad26 does not influence Pol II loading, but is required for an optimal induction of Pol II-transcribed genes. To further investigate Rad26 link to transcription, we examined whether, under standard growth conditions, any effect of *rad26* deletion on Mediator or Pol II occupancy could be identified. Our ChIP experiments did not show any difference between *rad26Δ* and WT strains in Mediator occupancy on selected UAS (**Supplementary Figure S4A**). Pol II occupancy of selected Pol II-transcribed genes in *rad26Δ* strain was similar to that of the wild-type (**Supplementary Figure S4B**). No growth phenotypes of *rad26Δ* were observed in GGR-proficient and deficient (*rad7Δ*) contexts (**Supplementary Figure S4C**), except UV sensitivity in *rad7Δ* context, as expected for this TCR component (**Supplementary Figure S1A**).

Taken together, our results show a strong correlation between Rad26 and Pol II on the yeast genome which is consistent with a close relationship between Rad26 and Pol II transcription, without Rad26 having a major role in this process. Moreover, this analysis revealed a colocalization of Rad26 and Mediator on intergenic regions.

### Genome-wide location analysis of Rad1-Rad10

Rad1-Rad10 endonuclease acts in NER as a dimer and together with Rad2 for the dual excision of damaged DNA. To determine whether these proteins are enriched on yeast chromatin in the absence of exogeneous genotoxic stress and whether their occupancies are correlated, we performed Rad1 and Rad10 ChIP-seq experiments. No major difference in UV sensitivity was observed for the strains carrying N-terminal or C-terminal HA-tagged Rad1 or Rad10 (**Supplementary Figure S1B**), suggesting that both HA-tagged versions were functional. Our ChIP experiments showed that N-terminally and C-terminally tagged Rad1 and Rad10 were enriched inside selected Pol II-transcribed genes, UASs, as well as Pol III-transcribed genes and telomeric regions (**Supplementary Figure S2B-D**), spanning similar groups of genomic targets as Rad2 (Eyboulet et al. 2013). The C-terminal tagged versions were used for ChIP-seq experiments, since the ChIP signal to noise ratios were higher (1.3-2 fold) compared to N-terminal tagged versions.

ChIP-seq analysis of Rad1 and Rad10 enrichment peaks revealed that they are distributed within Pol II-transcribed and intergenic regions, Pol III-transcribed genes, telomeric and centromeric regions (**Supplementary Figure S3**). Specific examples of ChIP-seq profiles illustrate Rad1 and Rad10 enrichment on Pol II-transcribed genes (**Figure 2A**). We compared genome-wide occupancy of Rad1 with that of Rad10 on Pol II-transcribed genes and observed a linear relationship on transcribed and intergenic regions (R^2^ = 0.94 and 0.82, respectively) that is consistent with the action of these two proteins as a dimer (**Figure 2B, C**). Spearman correlation analysis on Mediator enrichment peaks revealed a strong correlation between Rad1 and Rad10 sets of data (Spearman correlation coefficient equal to 0.79) and moderate correlation between Rad1/Rad10 and Mediator occupancies (Spearman correlation coefficient equal to 0.5 and 0.54, respectively) (**Figure 2D**). Heatmaps of tag density for Rad1, Rad10 and Mediator (Med17 subunit) on Mediator enrichment peaks, as well as metagene analysis demonstrated Rad1 and Rad10 enrichment on these regions (**Figure 2E**). In addition to Pol II-transcribed and intergenic regions, Rad1 and Rad10 were highly enriched on Pol III-transcribed genes showing a linear relationship (R^2^ = 0.94) (**Supplementary Figure S3C**), similar to previously observed enrichment of Rad2 on these genes (Eyboulet et al. 2013).

**Figure 2.**
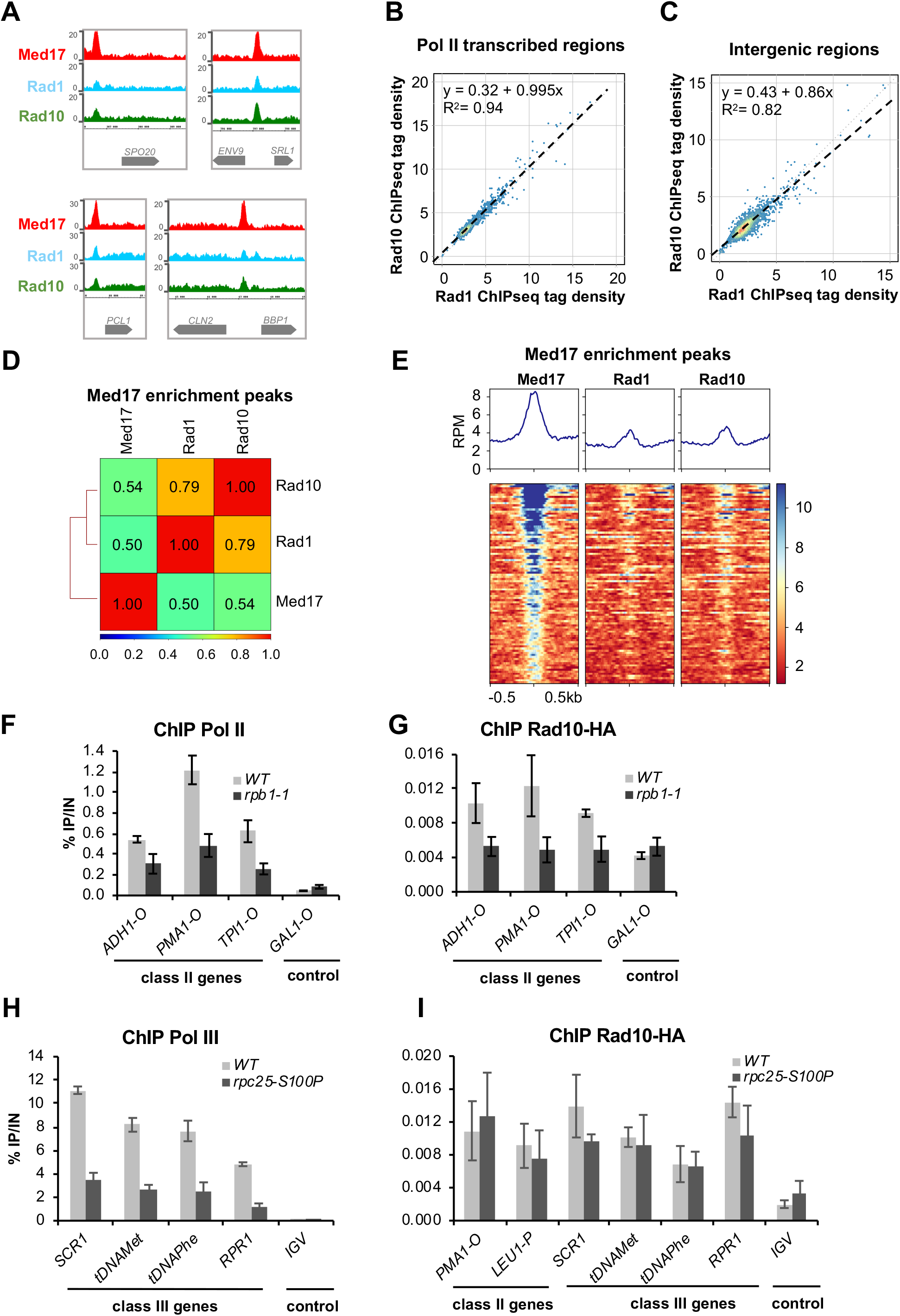
Genome-wide analysis of Rad1 and Rad10 occupancy. (**A**) Examples of Rad1, Rad10 and Mediator (Med17) tag density profiles in WT context on Pol II-transcribed and intergenic regions. (**B, C**) Rad10 ChIP-seq density versus Rad1 ChIP-seq density on Pol II transcribed regions (**B**) and intergenic regions (**C**). Each point on the plot corresponds to one transcribed or intergenic region. A linear regression (dotted line) and an R^2^ linear regression coefficient are indicated. The dashed line corresponds to y = x. **(D)** Pair-wise Spearman correlation coefficients (SCC) of ChIP-seq data were calculated for Rad1, Rad10 and Mediator on Mediator (Med17) enrichment peaks. The colors correspond to the scale for SCC indicated on the bottom. **(E)** Heatmaps of Mediator (Med17), Rad1 and Rad10 ChIP-seq profiles centered on Mediator (Med17) enrichment peaks (−500 bp to +500 bp), sorted by decreasing Mediator occupancy. Median tag density profiles in RPM (reads per million) are shown in upper panels. (**F, G**) Effect of *rpb1-1* Pol II mutation on Rad10 and Pol II occupancies on selected regions. Quantitative ChIP assays were performed using *α*-Rpb1 antibody (Pol II) (**F**), and *α*-HA antibody against Rad10-HA (**G**). Cells were grown in selective SD medium complemented with amino acids at 25°C and then shifted for 90 min at 37°C. *GAL1-O* amplicon was used as a negative control. Quantities were reported to qPCR performed on Input DNA and are expressed as a percentage. The indicated value is the mean of three biological replicates, and error bars represent the standard deviation. (**H, I**) Effect of *rpc25-S100P* Pol III mutation on Rad10 and Pol III occupancies on selected regions. Quantitative ChIP assays were performed using *α*-Myc antibody against Rpc160-13Myc Pol III subunit (**H**), and *α*-HA antibody against Rad10-HA (**I**). Cells were grown in selective SD medium complemented with amino acids at 25°C and then shifted for 10h at 37°C. *IGV* amplicon (non-transcribed region on chromosome V) was used as a negative control. Quantities were reported to qPCR performed on Input DNA and are expressed as a percentage. The indicated value is the mean of three biological replicates, and error bars represent the standard deviation.

To address the question of transcriptional dependency of Rad1 and Rad10 chromatin occupancy on Pol II-transcribed genes, we used a Pol II mutant, *rpb1-1*, that stops Pol II transcription after a shift to nonpermissive temperature (Nonet et al. 1987). As expected, Pol II occupancy was reduced on Pol II-transcribed (class II) genes in the *rpb1-1* mutant compared with the wild-type strain (**Figure 2F**). Rad10 occupancy on Pol II-transcribed genes followed that of Pol II and was also reduced in this mutant (**Figure 2G**). A decrease in Rad1 occupancy was also observed on selected Pol II-transcribed genes, even though the effect was less pronounced (**Supplementary S5A, B**). Thus, the presence of Rad1-Rad10 on Pol II-transcribed genes is dependent on Pol II transcription.

A Pol III-specific mutant, *rpc25-S100P*, was similarly used to impair Pol III transcription (Zaros and Thuriaux 2005) (**Figure 2H**). By contrast to Pol II-transcription dependency, no significant decrease was observed for Rad10 occupancy on Pol III-transcribed (class III) genes, suggesting that the presence of this protein on class III genes was independent on Pol III transcription (**Figure 2I**).

We tested whether *rad10* or *rad1* deletion leads to any growth phenotypes that could suggest a potential transcriptional implication for Rad1-Rad10. No growth difference between *rad10Δ* or *rad1Δ* and wild-type strains were observed under different temperature conditions, on different carbon sources or for NTP depletion conditions, except for a high UV sensitivity, as expected for key NER proteins (**Supplementary Figure S5C**). We also examined whether, under standard growth conditions, *rad10* or *rad1* deletion can affect Pol II and Mediator occupancy. We did not observe any differences between Pol II occupancy of the *rad1Δ* or *rad10Δ* strains compared to the wild-type strain on selected genes (**Supplementary Figure S5D, F**). Similarly, no effect of *rad1* or *rad10* deletion on Mediator occupancy on selected UASs was observed (**Supplementary Figure S5E, G**). Our results suggest that Rad1-Rad10 do not play a major role in Pol II transcription.

Taken together, our results show a strong correlation between Rad1 and Rad10 on the yeast genome, consistent with their function as a dimer, and revealed their presence on the chromatin without exogeneous genotoxic stress. Moreover, Rad1-Rad10 occupancy on Pol II-transcribed genes is transcription-dependent and they colocalize with Mediator on intergenic regions.

### Mediator interacts with Rad1/Rad10 endonuclease and Rad26 TCR-specific protein

Previously, we identified a physical link between Rad2 NER protein and Mediator complex that contributes to their functional interplay (Eyboulet et al. 2013). Given the co-localization of Rad26 and Rad1-Rad10 proteins with Mediator on the yeast genome, we examined their potential physical interactions with Mediator by CoIP experiments. Our results show that Mediator coimmunoprecipitates with Rad26, Rad1 and Rad10 in crude extracts of yeast strains expressing Med17-Myc and HA-Rad26, HA-Rad1 or Rad10-Flag (**Figure 3A-C**). No coimmunoprecipitation was observed between Mediator and two other NER proteins, Rad4 protein specific to GGR and Rad14 protein common for both NER pathways (**Supplementary Figure S6**). Interestingly, coimmunoprecipitation between Rad10 and Mediator remains unchanged in *rad2Δ* context, suggesting that Rad2 does not mediate this physical interaction (**Figure 3D**).

**Figure 3.**
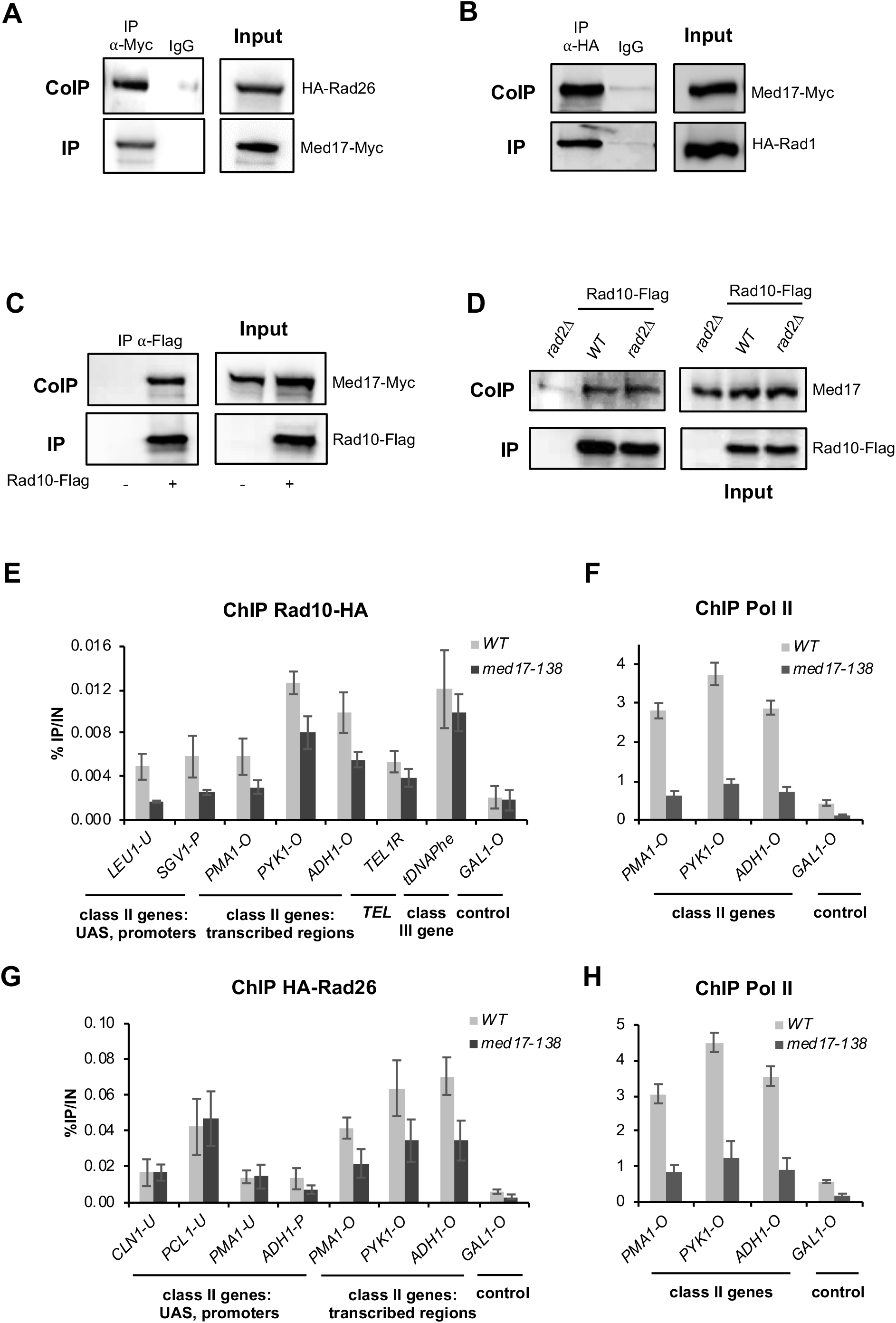
Rad1, Rad10 and Rad26 coimmunoprecipitate with Mediator and the loss of Mediator head in *med17-138* mutant decreases Rad10 chromatin binding. (**A-D**) CoIP between Rad1, Rad10 and Rad26 and Mediator. Inputs are shown on right panels. **(A)** Mediator was immunoprecipitated from crude yeast extracts via Med17-Myc subunit with *α*-Myc antibody (IP) and Western blotting with *α*-HA antibody detected HA-Rad26 (CoIP). **(B)** HA-Rad1 or (**C, D**) Rad10-Flag were immunoprecipitated with *α*-HA or *α*-Flag antibody, respectively, and analyzed by Western blotting with *α*-Myc antibody (**A-C**) against Med17-Myc subunit or rabbit polyclonal *α*-Med17 antibody (**D**) to detect Med17 Mediator subunit (CoIP) (left panels). WT (**A-D**) or *rad2Δ* (**D**) strains were used. IgG indicates a control immunoprecipitation with IgG magnetic beads only (**A, B**). A strain carrying non-tagged Rad10 was used as a negative control in **C** and **D**. (**E**-**H**) Effect of *med17-138* mutation on Rad10, Rad26 and Pol II occupancies on selected regions. Quantitative ChIP assays were performed using *α*-Rpb1 antibody (Pol II) (**F, H**), and *α*-HA antibody against Rad10-HA (**E**) or HA-Rad26 (**G**). Cells were grown in selective SD medium complemented with amino acids at 25°C and then shifted for 45 min at 37°C. *GAL1-O* amplicon was used as a negative control. Quantities were reported to qPCR performed on Input DNA and are expressed as a percentage. The indicated value is the mean of three biological replicates, and error bars represent the standard deviation.

To investigate whether the loss of the Mediator function influences Rad10 and Rad26 chromatin binding, we used *med17-138* mutant in which the Mediator head module dissociated from the rest of the complex at the non-permissive temperature (Thompson and Young 1995; Holstege et al. 1998; Linder et al. 2006; Takagi and Kornberg 2006). Our ChIP experiments show that Rad10 occupancy was reduced on most of the tested promoters, UASs and gene bodies of Pol II-transcribed genes in this Mediator mutant compared to the wild-type strain (**Figure 3E, Supplementary Figure S7A**). Conversely, Rad10 occupancy of telomeric and Pol III-transcribed regions did not change under the same conditions. As expected, Pol II occupancy was strongly reduced in *med17-138* (**Figure 3F**). We then examined how the *med17-138* mutation influences Rad26 chromatin association (**Figure 3G, Supplementary Figure S7B**). Our results demonstrated that, in this mutant, Rad26 occupancy on promoters and UASs of Pol II-transcribed genes remained similar to the wild-type strain. However, this Mediator mutation reduced Rad26 occupancy on Pol II-transcribed regions, together with a high decrease in Pol II occupancy, supporting a correlation between Rad26 and Pol II chromatin binding (**Figure 3G, H**).

Taken together, our results demonstrated that Rad1, Rad10 and Rad26 physically interact with the Mediator complex. Moreover, the loss of Mediator function led to a decrease in Rad10 occupancy on Pol II promoter, regulatory and transcribed regions, suggesting that Mediator influences the recruitment or stability of Rad10. On the contrary, no impact of the dissociation of the Mediator head on Rad26 presence on Pol II promoter or regulatory regions was observed. The presence of Rad26 was decreased only within gene bodies of Pol II-transcribed genes, accompanying reduced transcription.

### *Mediator stabilization in* kin28 *mutant impacts Rad1, Rad10, Rad26 chromatin binding*

To investigate how changes in Mediator and Pol II chromatin binding influence Rad1-Rad10 and Rad26 distribution on the yeast genome, we used a mutant in Kin28 TFIIH subunit, *kin28-ts16* (Cismowski et al. 1995). In this mutant, shifting to non-permissive temperature led to inhibition of Pol II CTD Ser5 phosphorylation. As a consequence of impaired Kin28 function, Mediator is stabilized on core promoters and Pol II is decreased on transcribed regions (Jeronimo and Robert 2014; Wong et al. 2014; Jeronimo et al. 2016; Petrenko et al. 2016). Previously, we used *kin28-ts* mutant to demonstrate that Mediator stabilization on core promoters led to major changes in Rad2 occupancy with a shift from UASs to core promoters and a decrease on Pol II-transcribed regions, showing the importance of dynamic interactions between Mediator, Rad2 and Pol II for Rad2 loading to the chromatin (Georges et al. 2019). To determine the genome-wide distribution of Mediator, Pol II, Rad26, Rad1 and Rad10 in *kin28* mutant compared to the wild-type, we performed ChIP-seq experiments for Med17 Myc-tagged Mediator subunit, Rpb1 Pol II subunit, HA-tagged Rad26, Rad1 and Rad10 after a shift to 37°C for 75 min. ChIP followed by qPCR was also performed on selected UAS, core promoters and transcribed regions (**Supplementary Figure S8**). In agreement with previous observation of Mediator stabilization on core promoters in *kin28* mutants (Jeronimo and Robert 2014; Wong et al. 2014; Jeronimo et al. 2016; Petrenko et al. 2016; Georges et al. 2019), metagene analysis and heatmaps centered on Med17 peaks in *kin28-ts* mutant (corresponding to core promoters) showed a large increase of Med17 occupancy in *kin28-ts* mutant compared to the wild-type, as well as a large decrease in Pol II occupancy (**Figure 4A**). Rad10 and Rad1 occupancies on Mediator peaks were increased in *kin28* mutant, even though, unlike the Mediator occupancy, no shift to core promoters can be observed for these proteins (**Figure 4B, Supplementary Figure S9**). The maximal Rad10 or Rad1 occupancy ratio between the mutant and the wild type was equal to 1.6 or 1.4, respectively (**Supplementary Figure S9**). Rad26 enrichment on Mediator peaks remained unchanged in *kin28* mutant compared to the wild-type. However, a large decrease of Rad26 occupancy was observed in *kin28* mutant on flanking regions correlated with a decrease in Pol II occupancy (**Figure 4B**). An average tag density analysis on intergenic regions shows a significant increase for Mediator, Rad1 and Rad10 occupancy in *kin28* strain compared to the WT (**Figure 4C**). Heatmaps centered on Mediator peaks corresponding to UASs as defined in the WT context illustrate these changes in Rad26, Rad10 and Rad1 occupancies that accompany the Mediator stabilization in *kin28* mutant (**Figure 4D**).

**Figure 4.**
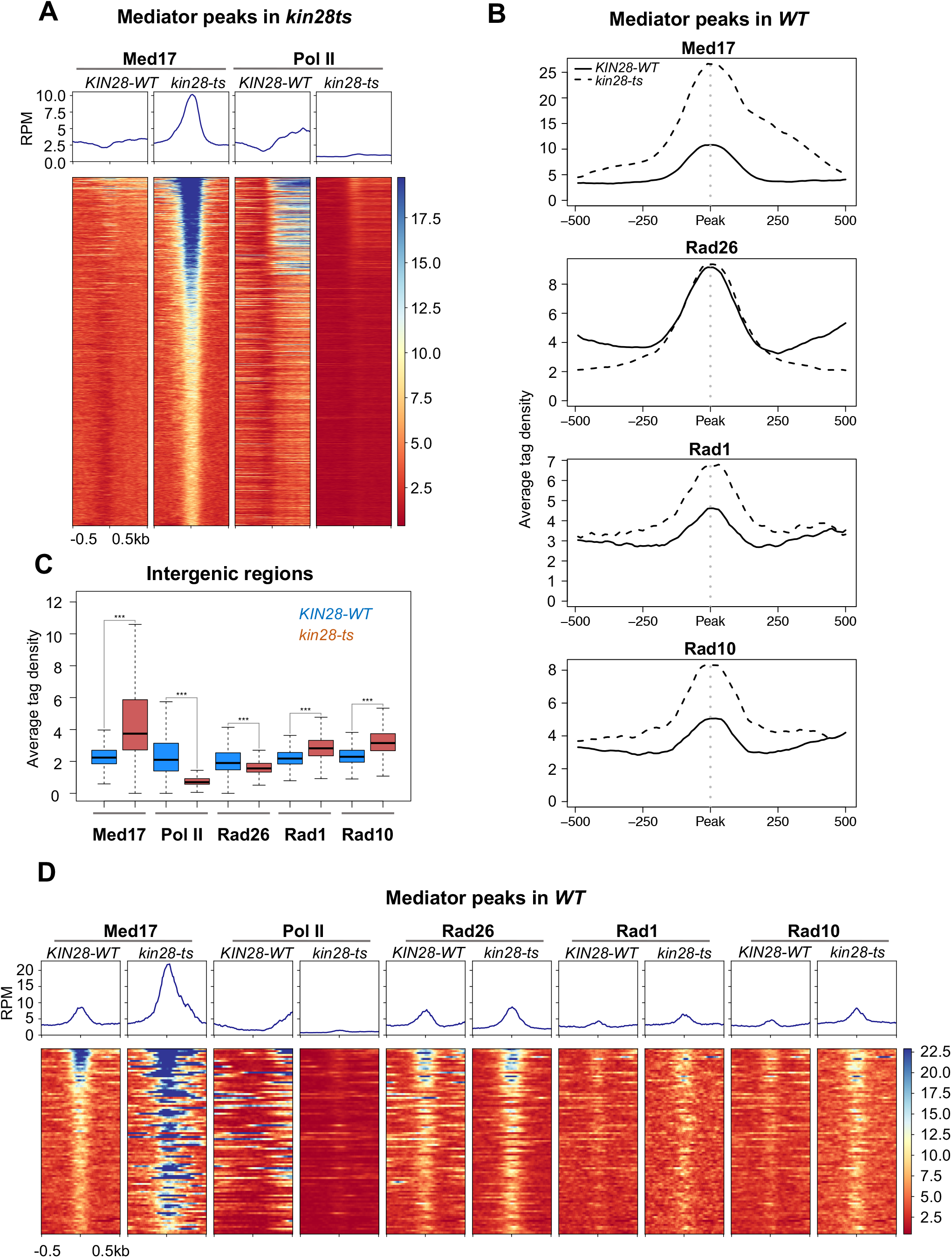
Effect of *kin28-ts* mutation on genome-wide Mediator, Rad26, Rad1 and Rad10 occupancy on intergenic regions. **(A)** Heatmaps of Mediator (Med17) and Pol II ChIP-seq profiles centered on Mediator (Med17) enrichment peaks determined in *kin28-ts* mutant (core promoters, -500 bp to +500 bp), sorted by decreasing Mediator occupancy. WT and *kin28-ts* strains were compared. Median tag density profiles in RPM (reads per million) are shown in upper panels. **(B)** Average tag density in Med17 Mediator, Rad26, Rad1 and Rad10 ChIP (from upper to lower panels), around Med17 Mediator enrichment peaks (−500 bp to +500 bp) determined in WT (U ASs). Average tag density in WT strains is indicated as a full line, whereas average tag density in *kin28-ts* strains is indicated as a dashed line. **(C)** Boxplots showing changes in Med17 (Mediator), Pol II, Rad26, Rad1 and Rad10 ChIP-seq on intergenic regions (intergenic regions for Pol II-transcribed genes in tandem or in divergent orientation, excluding intergenic regions encompassing Pol III-transcribed genes, centromeres, telomeres) in WT (in dark blue) and *kin28-ts* strains (in brown). The asterisks represent a significant difference between the WT and the mutant at p-value < 2.2e-16 in a Wilcoxon rank-sum test. **(D)** Heatmaps of Mediator (Med17), Pol II, Rad26, Rad1 and Rad10 ChIP-seq profiles centered on Mediator (Med17) enrichment peaks determined in WT (UASs, -500 bp to +500 bp), sorted by decreasing Mediator occupancy. WT and *kin28-ts* strains were compared. Median tag density profiles in RPM (reads per million) are shown in upper panels.

Rad26 occupancy was strongly reduced on transcribed regions, together with a strong decrease in Pol II occupancy in *kin28* mutant (**Figure 5A**). Rad10 and Rad1 occupancies also followed Pol II occupancy decrease. **Figure 5A** illustrates how *kin28* mutation affects Rad26, Rad10 and Rad1 occupancies, covering 10% of Pol II most-transcribed regions with 500bp before TSS and 500 bp after TES. Average tag density analysis on 10% Pol II most-enriched genes shows a large decrease in Pol II and Rad26 occupancies in *kin28* mutant, as well as a smaller but significant decrease in Rad1 and Rad10 (**Figure 5B**). Metagene analysis and heatmaps of ratios between *kin28* mutant and WT allow to visualize these changes in occupancy of Rad26, Mediator, Pol II, Rad1 and Rad10 (**Figure 5C, Supplementary Figure S10**).

**Figure 5.**
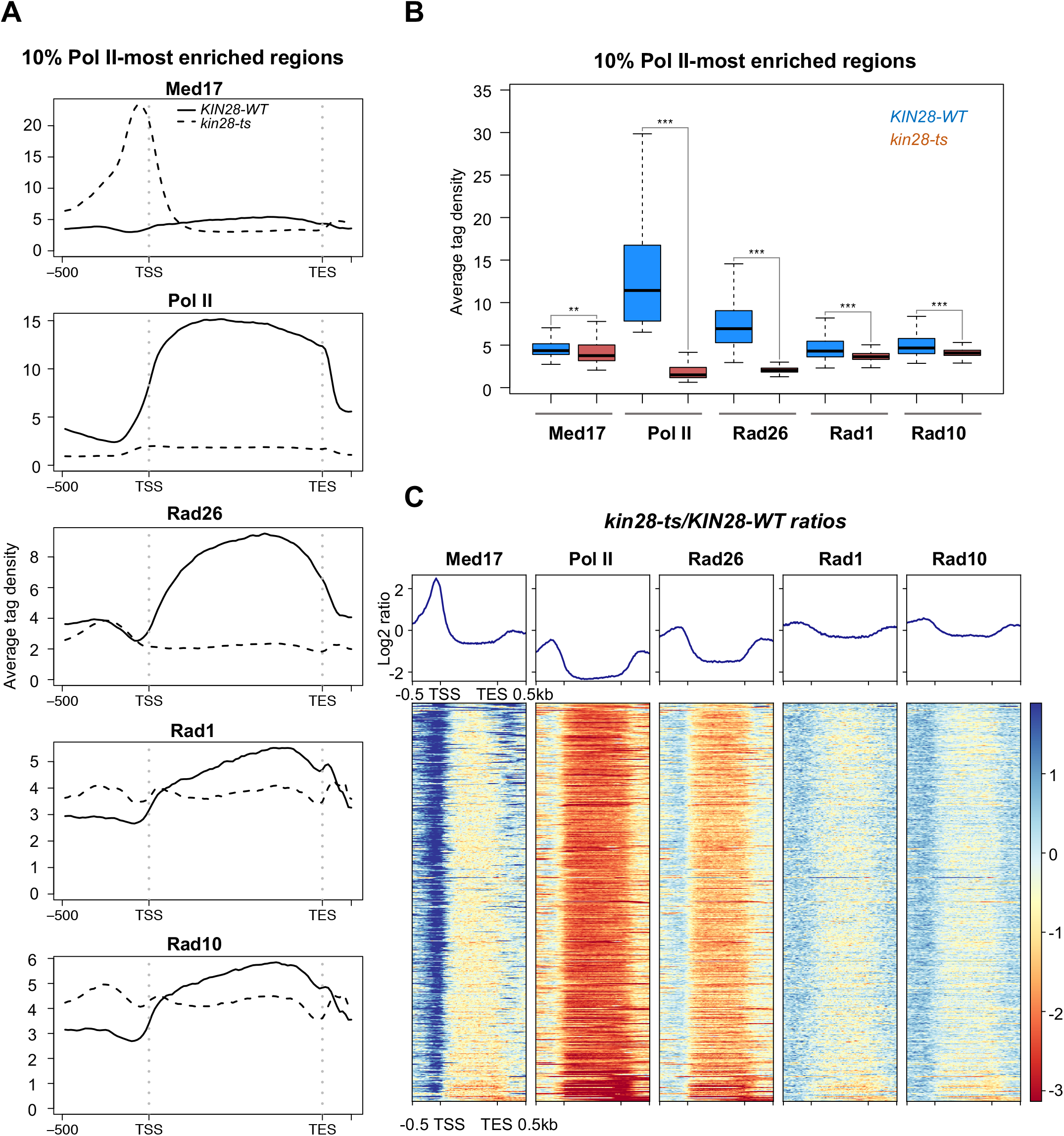
Effect of *kin28-ts* mutation on genome-wide Mediator, Pol II, Rad26, Rad1 and Rad10 occupancy on transcribed regions. **(A)** Average tag density in Med17 Mediator, Pol II, Rad26, Rad1 and Rad10 ChIP (from upper to lower panels) on 10% Pol II-most enriched regions (scaled windows for 500 bp before TSS, between TSS and TES, and 500 bp after TES). Average tag density in WT strains is indicated as a full line, whereas average tag density in *kin28-ts* strains is indicated as a dashed line. **(B)** Boxplots showing changes in Med17 (Mediator), Pol II, Rad26, Rad1 and Rad10 ChIP-seq on 10% Pol II-most enriched regions in WT (in dark blue) and *kin28-ts* strains (in brown). The asterisks represent a significant difference between the WT and the mutant at p-value < 1e-10 (**) or < 2.2e-16 (***) in a Wilcoxon rank-sum test. **(C)** Heatmaps of Mediator (Med17), Pol II, Rad26, Rad1 and Rad10 ChIP-seq occupancy ratios between *kin28-ts* mutant and WT on 10% Pol II-most enriched regions (scaled windows for 500 bp before TSS, between TSS and TES, and 500 bp after TES), sorted by decreasing Mediator occupancy ratio. Average log2 ratios are shown in upper panels.

Taken together, our results show that Rad1 and Rad10 chromatin binding in *kin28* mutant is influenced by Mediator that is stabilized on core promoters and by a decrease in Pol II transcription, whereas Rad26 occupancy in this mutant is mainly impacted by a large decrease in Pol II presence on gene bodies.

### Multivariate analysis of Mediator, Rad1, Rad10, Rad26 and Pol II genomic coverage signal on intergenic regions

To investigate in detail and find characteristics for complex relationship between different NER proteins and transcriptional components that co-occupy intergenic regions, we analyzed simultaneously the genome-wide data sets for Mediator, Pol II, Rad1, Rad10 and Rad26 in wild type and *kin28* mutant contexts. To this aim, we first applied the principal component analysis (PCA) for multivariate systems that allows to make a linear combination between system’s variables in order to evaluate the variability in data sets with a reduced number of parameters and construct an orthonormal coordinate system, where the orthogonal directions are statistically independent. Pareto representation showed that three principal components (PCs) are sufficient to explain 94.5% or 93.5% of the data variability in wt and *kin28* mutant, respectively (**Supplementary Figure S11A**). The space of three PCs (orthonormal referential {*PC*_1_, *PC*_2_, *PC*_3_}) was thus considered in further analysis (**Supplementary Figure S12)**. To understand how different variables (Mediator, Pol II, Rad1, Rad10 and Rad26) participate in the construction of PCs, the norm of each variable was calculated (**Supplementary Figure S11B**). In both contexts, Pol II and Mediator contribute the most to the observed data set variability, followed closely by Rad26, Rad1 and Rad10. We then measured the contribution of each variable to each PC (**Supplementary Figure S11C**). This analysis showed that Rad1, Rad10 and Rad26 proteins are the main contributors to PC1 in WT and *kin28* mutant. The PC2 is mainly influenced by Mediator and Pol II in a wild-type, while a *kin28* mutation changes the main contribution to Mediator and Rad1-Rad10. The three proteins Rad26, Rad1 and Rad10 are the main contributors to the definition of PC3 in WT, while Rad26 and Pol II influenced the most this PC in *kin28* mutant. Therefore, the stabilization of Mediator in *kin28* mutant affects mainly the interplay between Rad1-Rad10 and Rad26 proteins, and between Rad26 and Pol II.

To verify how variables covariate in the data set, we projected one variable over the other in the orthonormal reference of three PCs and calculated the covariance of the variables. In wild type context, we observed a strong correlation between Rad1 and Rad10 on intergenic regions (**Figure 6A, C**, third row – forth column), in accordance with the existence of a stable Rad1-Rad10 dimer. Pol II shows a positive correlation with Rad1, Rad10 and Rad26. However, while Rad26 shows a correlation with Pol II close to that with Mediator, this protein is not correlated with Rad10 and seems to be slightly anti-correlated with Rad1, suggesting that the link between Pol II and Rad26 could be in competition with that of Rad1-Rad10 dimer. The coverage of Mediator and Pol II are anti-correlated, in accordance with the Mediator occupancy on UASs located 200 to 500 bp upstream of TSS (Jeronimo and Robert 2014; Eyboulet et al. 2015; Paul et al. 2015) and Mediator shows a similar correlation with Rad26 and Rad10, suggesting that Mediator interplays with Rad26 and Rad10 independently of Pol II.

**Figure 6.**
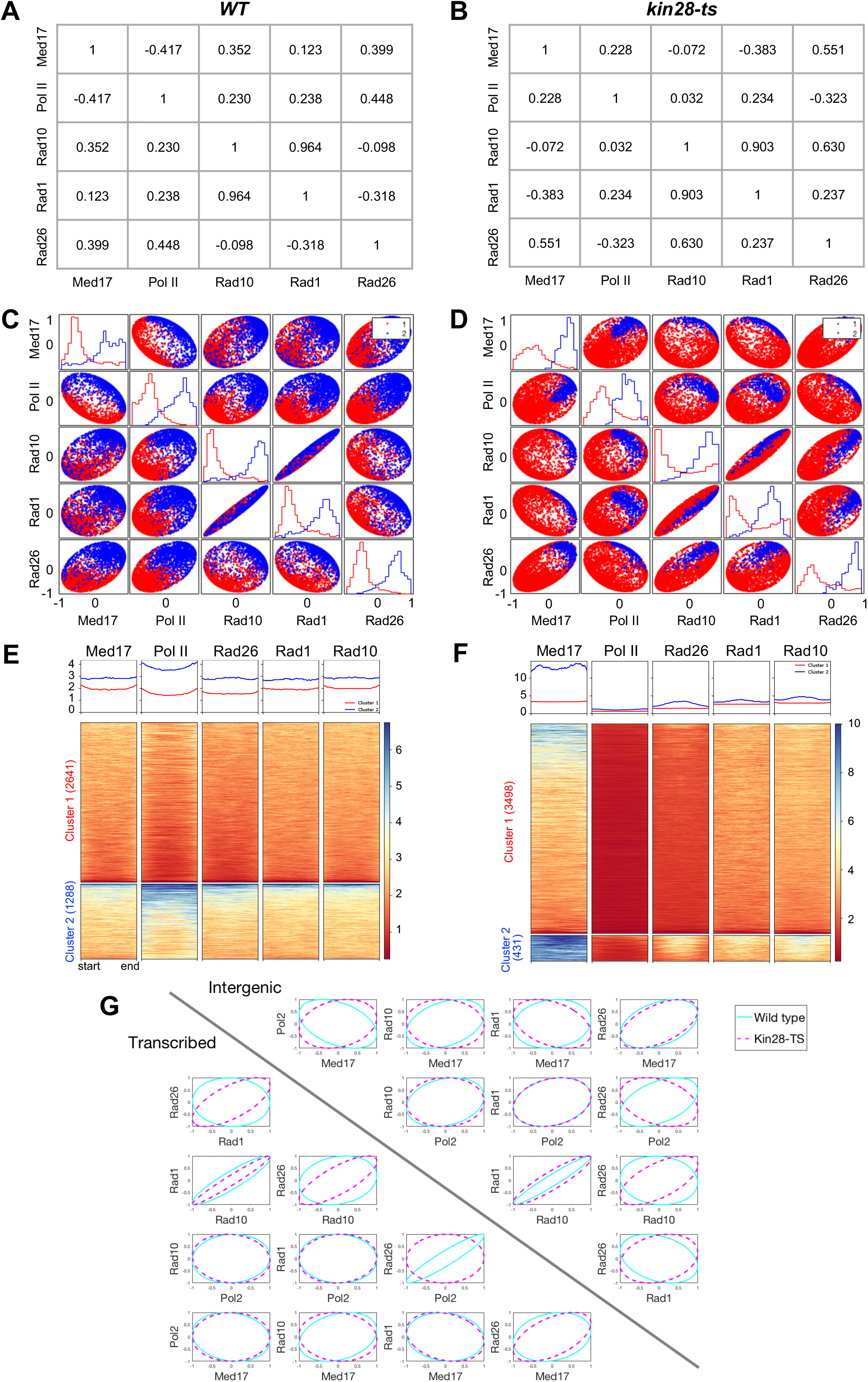
Multivariate and clustering analysis of relationship between Mediator, Rad1, Rad10, Rad26 and Pol II. (**A, B**) Covariance of the variables Mediator (Med17), Pol II, Rad1, Rad10 and Rad26 for ChIP-seq data on intergenic regions in WT (**A**) or *kin28-ts* (**B**) calculated as described in M&M. (**C, D**) Covariance distribution for ChIP-seq data on intergenic regions separated in two independent clusters in WT (**C**) or *kin28-ts* (**D**) context. The larger cluster 1 is represented in red and the smaller cluster 2 in blue. The diagonal represents the distribution of the covariance value for each cluster and for each variable. The width of the ellipsoid (small radius) corresponds to the dispersion of the relation between the two variables. (**E, F**) Heatmaps of Mediator (Med17), Pol II, Rad26, Rad1 and Rad10 ChIP-seq profiles on intergenic regions for clusters 1 and 2 in WT (**E**) and *kin28-ts* strains (**F**). Number of regions in each cluster is indicated in brackets. Median tag density profiles in RPM (reads per million) are shown in upper panels. (**G**) Comparison of covariance distribution for ChIP-seq data on intergenic (right upper panel) or transcribed regions (left lower panel) between WT (cyan contour) and *kin28* mutant (pink contour).

In *kin28* mutant, the strong correlation between Rad1 and Rad10 is maintained (**Figure 6B, D**). Interestingly, we observe a stronger correlation between Mediator and Rad26 (0.551 in *kin28* mutant versus 0.399 in WT). In this context, Rad26 becomes correlated with Rad10 and Rad1, and anticorrelated with Pol II, while the correlation between Rad1-Rad10 and Mediator is reduced (Rad10 loses its correlation with Mediator and Rad1 becomes anti-correlated). Therefore, the stabilization of Mediator on the chromatin in *kin28* mutant changes considerably the relationship between Mediator, Pol II and Rad26, Rad1 and Rad10, in particular it enhances the correlation of Mediator and Rad1-Rad10 with Rad26, and leads to an anti-correlation between Pol II and Rad26.

We then examined how homogeneous the data set behavior was by determining independent sub-ensembles within data points. To this purpose, we used the Silhouette plot (Rousseeuw 1987) on a Ward’s metrics obtained by a k-list hierarchical clustering method (Kaufman and Rousseeuw 1990). The Silhouette value was maximal for both wt and *kin28* mutant when the data sets were separated into two independent clusters (**Supplementary Figure S11D**). To understand how these two clusters are distributed in terms of covariance, we plotted the covariance between the variables and used a color code to represent each cluster (**Figures 6C, D**). The diagonal represents the distribution of the covariance values for each cluster and each variable in PCs referential. This ellipsoid representation allows to visualize correlation or anti-correlation relationship between variables. The width of the ellipsoid (small radius) corresponds to the dispersion of the relation between the two variables. Interestingly, we observed that in the wild type the cluster 1 (red dots) corresponds to data points that are oriented in the opposite directions (negative values) and the cluster 2 (blue dots) corresponds to data points that are co-oriented (positive values) with the variable direction (**Figures 6C)**. In other words, the oppositely orientated data points correspond to those data points that have the smallest contribution to the definition of the variable direction and the positive ones are those that define the direction of the variables. This interpretation is confirmed by the heatmap of the two defined clusters, where the elements of the second cluster have higher occupancy than the elements of the first cluster (**Figures 6E, F**).

The *kin28* mutation increases dispersion between variables, that is particularly well observed by the increase in the size of the small radius of the ellipsoid representing covariance distribution between Rad1 and Rad10 (**Figures 6D**). Interestingly, the size of the cluster 1 is increased and the size of the cluster 2 is reduced in the mutant context (**Figures 6E, F**), meaning that the stabilization of Mediator in *kin28* mutant leads to a homogeneous level of coverage of the other proteins on intergenic regions, hence reducing the number of regions with higher-than-average coverage values. This observation suggests that the stabilization of Mediator on the chromatin modifies the relationship between Mediator, Rad1-Rad10, Rad26 and Pol II. Importantly, *kin28* mutation increases a correlation between Mediator and Rad26 or Rad1-Rad10 and Rad26.

To see how two clusters are distributed for the tag density of different proteins, we generated heatmaps in wt and *kin28* mutant (**Figure 6E, F**). In wild-type context, the cluster 2 corresponding to one third of regions is characterized by higher occupancy of all analyzed proteins compared to the cluster 1 (**Figure 6E**). In *kin28* mutant, the cluster 1 represents a majority of regions and the cluster 2 encompassing 431 regions is defined by the highest enrichment of Mediator (**Figure 6F**). This cluster 2 highly-enriched by Mediator is also more enriched by Rad26, Rad1 and Rad10. **Supplementary Figure S13** shows an overlapping between clusters in WT and *kin28* mutant.

To illustrate how *kin28* mutation changes the relationship between variables, ellipsoid contours of covariance distribution were compared between wt (**Figure 6C)** and *kin28* mutant (**Figure 6D)** on the intergenic and transcribed regions by plotting them on the same graph (**Figure 6G**, right upper and left lower triangles, respectively). This comparison on intergenic regions (right upper triangle) clearly shows that covariance distribution between Pol II and Rad10 or Rad1, or between Rad26 and Mediator did not change the orientation in *kin28* mutant compared to the wild-type, whereas *kin28* mutation leads to an opposite orientation between Rad26 and Pol II, Rad1 or Rad10. For comparison, we also performed the same analysis on transcribed regions (left lower triangle) and showed that in *kin28* mutant a correlation between Rad26 and Rad1 or Rad10 was increased, whereas a strong correlation between Rad26 and Pol II was lost.

In conclusion, by taking advantage of the orthonormal referential that one can construct using Principal Component analysis, we formulated a data representation that shows the simultaneous relations among coverage of chromatin by different interacting proteins. This representation allowed us to reveal the interplay between Rad1, Rad10 and Rad26 proteins, Mediator and Pol II and their regulation by the binding dynamic of Mediator.

## Discussion

In this study, we performed genome-wide location analysis of Rad1/XPF, Rad10/ERCC1 and Rad26/CSB NER proteins in yeast *Saccharomyces cerevisiae*, addressing the questions on their chromatin binding in the absence of exogeneous genotoxins and their potential interplay with Mediator and Pol II. Our results showed that Rad26 and Rad1-Rad10 associate to a wide range of regions in the absence of exogeneous genotoxic stress, indicating that the chromatin binding of these proteins is not restricted to DNA damage-inducing conditions. However, despite similarities in genomic localization, we demonstrated that their functional link with transcription, Pol II and Mediator differs substantially, suggesting different modes of recruitment and/or function.

NER being coupled with transcription, we investigated the relationship between Rad1-Rad10, Rad26 and Pol II. Consistent with previous studies, Rad26 is the most closely related to Pol II transcription. All the proteins show positive correlation with Pol II, with Rad26 being very highly correlated (**Figure 1B**, SCC=0.91). In the *rpb1-1* Pol II mutant, the chromatin association of all three NER proteins decreased ((Malik et al. 2010), **Figure 2G, Supplementary Figure S5B**), showing that their presence on Pol II-transcribed genes is transcription-dependent. In *kin28* TFIIH mutant with a strong decrease in Pol II transcription, Rad26 occupancy was strongly reduced on transcribed regions and Rad1-Rad10 occupancies also followed this decrease (**Figure 5A**). However, Pol II occupancy does not change if either of Rad1, Rad10 or Rad26-coding genes are deleted (**Supplementary Figures S4B, S5D, F**), suggesting that Pol II transcription does not absolutely require these proteins. Moreover, a deletion of *rad1, rad10* or *rad26* does not lead to any growth phenotypes, with an exception of UV sensitivity (in GGR-deficient background for *rad26*), providing further arguments that these proteins do not play a major role in Pol II transcription. Thus, the loading of Rad1, Rad10 and Rad26 proteins on chromatin could physically depend on Pol II presence and/or could be required to resolve potential genomic instability associated with Pol II transcriptional activity. It should be also noted that Rad26/CSB has a DNA-dependent ATPase activity that can be important for chromatin remodeling and regulation of chromatin structure (Selby and Sancar 1997b; Newman et al. 2006; Lake et al. 2014), helping Pol II to pass through nucleosomes (Xu et al. 2020) and leading to Rad26 requirement for TCR downstream of the +1 nucleosome (Duan et al. 2020).

We previously discovered the link between Rad2 and Mediator, showing that Rad2 chromatin binding depends on Pol II transcription and involves dynamic interactions with Mediator and Pol II (Eyboulet et al. 2013; Georges et al. 2019). In this work, we showed that Mediator link to NER proteins is not restricted to Rad2, but it also includes Rad1-Rad10 and Rad26. Mediator colocalizes with Rad1, Rad10 and Rad26 on the yeast genome (**Figure 1D, 2E**) and physically interacts with these proteins (**Figure 3A-D**). Moreover, changes in Mediator function affect NER factors, although to a different extent. The genome-wide co-occupancy of Mediator-enriched regulatory regions and physical interactions between Mediator and Rad1, Rad10 and Rad26 NER proteins were similar to previous results for Rad2 (Eyboulet et al. 2013). Surprisingly, considerable differences were observed between Rad1-Rad10 and Rad26 with respect to changes in Mediator function and chromatin binding. The loss of Mediator function induced by the dissociation of the Mediator head in *med17-138* mutant led to a significant decrease in Rad10 occupancy on regulatory, promoter and transcribed regions of Pol II-dependent genes (**Figure 3E**). This result suggests that Mediator and in particular Mediator head module has an impact on Rad10 recruitment or stability on yeast genome. The situation was rather different for Rad26: no changes were observed for occupancy of this protein on regulatory and promoter regions in *med17-138* mutant compared to the wild type, whereas the presence of Rad26 on transcribed regions decreased, accompanying Pol II transcription deficiency (**Figure 3G**). This suggests that Rad26 recruitment on regulatory and promoter regions was independent of the Mediator head module. We could not exclude the possibility that other Mediator modules influence Rad26 recruitment. Moreover, when Mediator was stabilized on core promoters in *kin28* TFIIH mutant, Rad1 and Rad10 occupancy increased on regulatory regions, whereas Rad26 occupancy of these regions remained similar to the wild type (**Figure 4B**). In both Mediator and TFIIH mutants, Rad1-Rad10 were more impacted by changes in Mediator function or binding than Rad26. Similar to Pol II relationship, deletion of either of the genes led to unchanged Mediator binding levels (**Supplementary Figures S4A, S5E, G**). Among NER factors, the Rad2 genomic binding shows the highest correlation with Mediator on regulatory regions (Eyboulet et al. 2013), as well as a stabilization and a shift to core promoters accompanying Mediator stabilization on core promoters in *kin28* mutant (Georges et al. 2019). Our results show that Rad1 and Rad10 presence is also influenced by Mediator though to a lesser extent, whereas Rad26 is the least impacted.

Using a multivariate method, we classified intergenic and transcribed regions of the genome with respect to Mediator, Pol II, Rad1, Rad10 and Rad26 protein coverage. Our genome-wide analyses uncover a high correlation between Rad1 and Rad10 proteins, consistent with their function as a dimer. We found that in wild type context while the majority of the regions have a comparable level of protein coverage, some regions have higher protein coverage level. Notably, this analysis reveals that in wild type context Rad26 and Rad1-Rad10 dimer coverages are anti-correlated, but both coverages are correlated with that of Pol II, suggesting that Rad26 and Rad1-Rad10 could be competing to interplay with Pol II. In the same manner while Rad1 coverage is not correlated with that of Mediator, Rad 10 coverage is correlated, suggesting that Rad1-Rad10 link with Mediator could be mediated by Rad10. The same analysis in *kin28* TFIIH mutant showed that the number of regions with higher level of protein coverage is reduced, in agreement with the stabilization of Mediator changing the interplay between Pol II, Rad1-Rad10, Rad26 and Mediator. However, our analysis can fine tune this idea and suggests that while the stabilization of Mediator disrupts the interplay between Rad26 and Pol II, it enhances the relationship between Rad26 and Rad1-Rad10 dimer or Mediator.

We showed that in addition to Pol II-transcribed genes and intergenic regions, Rad1, Rad10 and Rad26 are also enriched on Pol III-transcribed genes, telomeres and centromeres, opening interesting perspectives for possible functions of these proteins. Rad2 was also present on Pol III-transcribed genes, telomeres and centromeres ((Eyboulet et al. 2013) and additional analysis). A complex genomic distribution of these proteins and their colocalization on a number of genomic features could be related to their different functions that remain to be fully uncovered. Thus, in addition to their role in NER, Rad1-Rad10/XPF-ERCC1 are also involved in other DNA repair mechanisms including DNA inter-strand crosslink (ICL) and DNA double-strand break (DSB) repair. Participation of CSB in repair of DSB and oxidative DNA damages (Charlet-Berguerand et al. 2006; Batenburg et al. 2015; Wei et al. 2015; Menoni et al. 2018; Pascucci et al. 2018), and in transcription-coupled homologous recombination (Teng et al. 2018) was proposed. A role of XPF-ERCC1 and CSB was also suggested in telomere maintenance (Zhu et al. 2003; Vannier et al. 2009).

In addition to DNA repair, XPF together with XPG/Rad2 and CSB were proposed to promote R-loop-induced genome instability (Sollier et al. 2014). Interestingly, genome-wide analysis of R-loops on yeast genome pointed out a particular enrichment of these structures on Pol III-transcribed genes and telomeres (Chan et al. 2014; El Hage et al. 2014; Santos-Pereira and Aguilera 2015; Wahba et al. 2016). It has been also suggested that R-loops are formed at centromeres and pericentromeric regions and that they are linked to histone H3 S10 phosphorylation and chromatin condensation (Castellano-Pozo et al. 2013). The enrichment of Rad2, Rad1-Rad10 and Rad26 on genomic regions predisposed to form R-loops including Pol III-transcribed genes, telomeres and centromeres raises a possibility that the recruitment of these proteins is linked to RNA-DNA hybrid formation or resolution. On centromeric and peri-centromeric regions, we noted that Rad1, Rad10 and Rad26 distributions follow a same tendency with a few highly-enriched centromeres and a bimodal profile for the remaining centromeres enriched on pericentromeric regions, requiring further investigation to propose a reason of this particular distribution. Interestingly, Rad26 and Rad1-Rad10 enrichment profiles on Pol II-transcribed genes with a higher signal on 3’-end (**Figure 5A**) resemble those observed for R-loops, in comparison with more equal distribution of Pol II along the genes (Wahba et al. 2016). For intron-containing genes, this R-loop profile was proposed to be related to higher enrichment of second exons (El Hage et al. 2014).

In conclusion, our genome-wide location analysis reveals that Rad1-Rad10 and Rad26 NER proteins are present on the yeast chromatin in the absence of exogeneous genotoxic stress, raising important questions and providing interesting perspectives for future research. Our results suggest that Mediator link to NER is not restricted to Rad2 protein and can be more complex, providing new information on functional dynamics between Rad1-Rad10, Rad26, Pol II and Mediator on the genomic scale.

## Materials and Methods

### Strains

All *S. cerevisiae* strains used in this study can be found in **Supplementary Table S1**.

### Spotting assay

Yeast cells were grown on plates at 30°C for 3 days. Cells were then scraped and diluted in water at 1 OD_600_. The yeast suspension was serially diluted by 10-fold and 3 μL was spotted on different media (YP medium supplemented with glucose, galactose, ethanol, or glycerol) or UV-irradiated at 5 or 20J/m^2^ (UV Stratalinker 1800) and incubated for 3 days or 5 days for Mycophenolic acid (25 or 50 μg/mL). Cells were grown at 30°C or at indicated temperature.

### Co-IP experiments

Co-immunoprecipitation experiments were conducted as previously described (Georges et al. 2019). Briefly, cells were grown at 30°C and 100 mL of exponentially grown cells were collected by centrifugation. Protein extracts were obtained by bead-beating in WB buffer (10% Glycerol, 50mM Hepes-KOH pH 7.5, 150mM NaCl, 1mM ethylenediaminetetraacetic acid (EDTA), 0.05% NP-40, 1 mM dithiothreitol (DTT), cOmplete protease inhibitor cocktails (Roche), 1 mM PMSF).

50 µL of Dynabeads pan-mouse IgG were incubated for 1h at 4°C with antibodies (70 ng/μL anti-HA (12CA5), 30 ng/μL anti-Myc (9E10) or 80 ng/μL anti-Flag (M2)). Antibody coated beads were washed three times with 0.1% BSA-PBS and twice with the WB buffer. The beads were then incubated with 1.5 mg of protein extract for 3h at 4°C. The beads were then washed four times with WB buffer. The beads and the input samples were resuspended in 40μL Novex™ Tris-Glycine SDS Sample Buffer (diluted to 1X) supplemented with 10% DTT 1M. The samples were kept at -80°C until further analysis.

Beads were heated at 85°C for 5min before performing gel electrophoresis. Separation was done on 8% or 12% bis-acrylamide gels in Tris-Glycine-SDS buffer. Proteins were then transferred on Amersham Protran 0.2 NC membranes (GE Healthcare). Membranes were blocked overnight in Tris-buffered-saline supplemented with 0.5% Tween 20 (TBS-T) and 5% milk. Membranes were incubated 1 h with the corresponding primary antibody in TBS-T with 2% milk (diluted at 1:10000 for anti-HA (12CA5), or 1:5000 for anti-Myc (9E10)). Membranes were washed three times in TBS-T before incubating with secondary antibodies (HRP-anti Mouse-IgG (H+L) (Promega)) for 45 min in TBS-T and 2% milk. After three more washes in TBS-T, detection was carried out using Amersham ECL or ECL-Prime reagents (GE Healthcare). Imaging was done using a Fusion FX7 imaging system.

### ChIP experiments

ChIP experiments were performed in triplicates. 100 mL of three independent cultures were grown as indicated below, depending on the strain used.

- *rad2, rad26, rad1* and *rad10* deletion mutants, and the corresponding wild-types were grown at 30°C.
- In *RPB1 wild-type* and *rpb1-1* mutant contexts, cells were grown at 25°C to mid-exponential phase and then shifted to 37°C for 90 min.
- In *RPC25 wild-type* and *rpc25-S100P* mutant contexts, cells were grown at 25°C to mid-exponential phase and then shifted to 37°C for 10 h.
- In *MED17* and *med17-138* mutant contexts, cells were grown at 25°C to mid-exponential phase and then shifted to 37°C for 45 min.
- In *KIN28 wild-type* and *kin28* mutant contexts, cells were grown at 25°C to mid-exponential phase and then shifted to 37°C for 75 min.

ChIP experiments were performed as follows, except for *kin28-ts* and the corresponding WT samples which were done on an IP-Star compact automated system (Diagenode) as previously described in (Georges et al. 2019). Cells exponentially growing (0.6 OD_600_) in 100 mL of YPD were treated with formaldehyde at a final concentration of 1% for 10 min for DNA-protein cross-linking. The reaction was stopped by adding glycine to a final concentration of 500 mM and incubating for 5 min. Cells were washed twice with cold 20 mM Tris-HCl pH8. Cells were then lysed by bead-beating for 30 min at 4°C in a FA/SDS buffer (50 mM Hepes KOH pH 7.5, 150 mM NaCl, 1 mM EDTA, 1% Triton-100, 0.1% sodium deoxycholate, 0.1% SDS) containing PMSF (FA/SDS+PMSF). Chromatin was sheared using a S220 focused-ultrasonicator (Covaris) for 10 min at 150W and duty factor 10, to obtain DNA fragments of about 200 bp.

50 µL of Dynabeads® Pan Mouse IgG was washed twice with cold PBS and then incubated with anti-HA (12CA5, 70 ng/μL), anti-Myc (9E10, 30 ng/μL) or anti-Rpb1-CTD (8WG16100 ng/μl) in cold PBS-0.1% BSA for 1h at 4°C. The antibody-coated beads were washed twice with cold PBS-0.1% BSA for 10min and one quick wash with FA/SDS buffer. Sonicated chromatin (about one-fourth of the total) and antibody-coated beads were incubated for 2h at 21°C under agitation. Beads were washed thrice with FA/SDS + 500 mM NaCl. The beads were then washed with IP Buffer (10mM Tris-HCl pH8, 0.25M LiCl, 1mM EDTA, 0.5% NP40, 0.5% sodium deoxycholate), followed by one wash in TE buffer (10mM Tris-HCl pH 8, 1mM EDTA). Elution was carried out at 65° C for 20 min, under agitation in a pronase buffer. Pronase, at a final concentration of 1mg/mL, was added to the eluate and incubated at 37 °C for 30 min. For the input, sample was also treated with pronase. Cross-links were reversed by overnight incubation at 65°C. RNase A at a final concentration of 0.025mg/mL was added to immunoprcipitated DNA (IP) and Input DNA (IN), and incubated at 37 °C for 1h. DNA was purified using Qiagen kit for PCR purification, according to manufacturer’s protocol. Three independent biological replicates were used for quantitative analysis. qPCR experiments were carried out using qPCR MasterMix SYBR® Green (Taykon) and primers listed in **Supplementary Table S2**.

For ChIP-seq, *KIN28 WT* and *kin28-ts* mutant were grown at 25°C to mid-exponential phase and then shifted to 37°C for 75 min. Three independent biological replicates were combined and library preparation for sequencing was made as described in (Georges et al. 2019).

### Data analysis

ChIP-seq data were analyzed using the following procedure. Reads were first trimmed with cutadapt (v1.12, http://dx.doi.org/10.14806/ej.17.1.200) then mapped on *S. cerevisiae* genome (University of California at Santa Cruz [UCSC] version sacCer3) using bowtie2 (v2.3.4.3, (Langmead and Salzberg 2012)). Files were converted using Samtools (v0.1.18, (Li et al. 2009)) and deepTools (v3.3.0, (Ramirez et al. 2016)), only unique reads were kept. Read counts were first normalized in RPM (Reads Per Million of mapped reads) then by qPCR data, on a set of selected regions, using the ratio between WT and mutant strains as previously described (Eyboulet et al. 2015). The number of mapped reads for each ChIP-seq experiment and normalization coefficients are indicated in **Supplementary Table S3**.

The transcribed (mRNA) regions were determined using the TSS (Transcription Start Sites) and TES (Transcription End Sites) of mRNA genes taken from (Pelechano et al. 2013; Malabat et al. 2015) (n=5337). ARS (n=196), Centromeres (n=16), telomeres (n=32), snRNAs (n=6), snoRNAs (n=77), tRNAs (n=299), rDNA (n=27), LTRs (n=383), and retrotransposons (n=50) coordinates were taken from yeastmine database https://doi.org/10.1093/database/bar062]. Intergenic regions were defined as non-transcribed regions not overlapping any of the previously listed genome features or their direct flanking sequences, resulting in 3929 regions. Peaks were detected with MACS2 (v2.1.3), using the input dataset as control (Zhang et al. 2008). Only peaks passing the quality filter were kept (fold_change>=1.5x and p-value<1e-10). Genome distribution of the peaks was determined based on their summit coordinates. For the heatmaps, peaks were oriented according to the closest TSS (maximum 1kb away) and only peaks located in the intergenic regions were used.

Heatmaps and correlation matrices were generated by deeptools. R (v3.5.1, https://www.R-project.org/) was used to generate profiles, boxplots, dotplots (ggplot2, (Wickham 2009)). The asterisks in boxplots represent a significant difference between the WT and the mutant at p-value < 0.05 (*) or < 1e-10 (**) or < 2.2e-16 (***) in a Wilcoxon rank-sum test [http://doi.org/10.1080/01621459.1972.10481279].

PCA and clustering were performed as follows. The number of PCs used in PCA was determined by Pareto chart of a distribution of sample variance as a function of PCs that allows to visualize the % of explained total variance by PCs (Wilkinson 2006). To evaluate how different variables participate in the construction of PCs, the norm of each variable in the orthonormal referential defined by {*PC*_1_, *PC*_2_, *PC*_3_} was calculated as 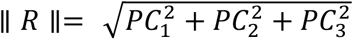. Larger is the norm value, more the variable contributes to the variability observed in a data set. The contribution of each variable to three PCs was expressed in the normal referential {*Med* 17, *Pol* II, *Rad* 1, *Rad* 10, *Rad* 26}. To verify how variables covariate in the data set, one variable was projected over the other in the orthonormal referential {*PC*_1_, *PC*_2_, *PC*_3_} as 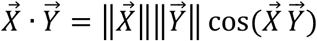. The covariance of the variables was calculated by normalizing the obtained product by the vector norms. For clustering analysis, we used the Silhouette Algorithm on a Ward’s metrics obtained by a k-list hierarchical clustering method (Rousseeuw 1987; Kaufman and Rousseeuw 1990). Higher is the Silhouette value, better the data set is separated into corresponding number of clusters. The covariance between variables was plotted using an ellipsoid representation with the width of the ellipsoid (small radius) corresponding to the dispersion between the two variables. The distribution of the covariance value for each cluster and for each variable was presented on the diagonal.

## Data availability

The ChIP-seq data have been deposited to the Array Express under the accession number E-MTAB-10828.

## Acknowledgements

We acknowledge the High-throughput sequencing facility of I2BC for its sequencing and bioinformatics expertise, the SPI (CEA/Saclay) for monoclonal antibodies, M. Werner and S. Marcand for fruitful discussions.

## References

Bailly V, Sommers CH, Sung P, Prakash L, Prakash S. 1992. Specific complex formation between proteins encoded by the yeast DNA repair and recombination genes RAD1 and RAD10. Proc Natl Acad Sci U S A 89: 8273–8277.

Batenburg NL, Thompson EL, Hendrickson EA, Zhu XD. 2015. Cockayne syndrome group B protein regulates DNA double-strand break repair and checkpoint activation. EMBO J 34: 1399–1416.

Boetefuer EL, Lake RJ, Fan HY. 2018. Mechanistic insights into the regulation of transcription and transcription-coupled DNA repair by Cockayne syndrome protein B. Nucleic Acids Res 46: 7471–7479.

Bukowska B, Karwowski BT. 2018. Actual state of knowledge in the field of diseases related with defective nucleotide excision repair. Life Sci 195: 6–18.

Castellano-Pozo M, Santos-Pereira JM, Rondon AG, Barroso S, Andujar E, Perez-Alegre M, Garcia-Muse T, Aguilera A. 2013. R loops are linked to histone H3 S10 phosphorylation and chromatin condensation. Molecular cell 52: 583–590.

Chan YA, Aristizabal MJ, Lu PY, Luo Z, Hamza A, Kobor MS, Stirling PC, Hieter P. 2014. Genome-wide profiling of yeast DNA:RNA hybrid prone sites with DRIP-chip. PLoS Genet 10: e1004288.

Charlet-Berguerand N, Feuerhahn S, Kong SE, Ziserman H, Conaway JW, Conaway R, Egly JM. 2006. RNA polymerase II bypass of oxidative DNA damage is regulated by transcription elongation factors. EMBO J 25: 5481–5491.

Cismowski MJ, Laff GM, Solomon MJ, Reed SI. 1995. KIN28 encodes a C-terminal domain kinase that controls mRNA transcription in Saccharomyces cerevisiae but lacks cyclin-dependent kinase-activating kinase (CAK) activity. Mol Cell Biol 15: 2983–2992.

Cleaver JE, Lam ET, Revet I. 2009. Disorders of nucleotide excision repair: the genetic and molecular basis of heterogeneity. Nat Rev Genet 10: 756–768.

Duan M, Selvam K, Wyrick JJ, Mao P. 2020. Genome-wide role of Rad26 in promoting transcription-coupled nucleotide excision repair in yeast chromatin. Proc Natl Acad Sci U S A 117: 18608–18616.

El Hage A, Webb S, Kerr A, Tollervey D. 2014. Genome-wide distribution of RNA-DNA hybrids identifies RNase H targets in tRNA genes, retrotransposons and mitochondria. PLoS Genet 10: e1004716.

Epanchintsev A, Costanzo F, Rauschendorf MA, Caputo M, Ye T, Donnio LM, Proietti-de-Santis L, Coin F, Laugel V, Egly JM. 2017. Cockayne’s Syndrome A and B Proteins Regulate Transcription Arrest after Genotoxic Stress by Promoting ATF3 Degradation. Molecular cell 68: 1054–1066 e1056.

Eyboulet F, Cibot C, Eychenne T, Neil H, Alibert O, Werner M, Soutourina J. 2013. Mediator links transcription and DNA repair by facilitating Rad2/XPG recruitment. Genes & development 27: 2549–2562.

Eyboulet F, Wydau-Dematteis S, Eychenne T, Alibert O, Neil H, Boschiero C, Nevers MC, Volland H, Cornu D, Redeker V et al. 2015. Mediator independently orchestrates multiple steps of preinitiation complex assembly in vivo. Nucleic Acids Res 43: 9214–9231.

Faridounnia M, Folkers GE, Boelens R. 2018. Function and Interactions of ERCC1-XPF in DNA Damage Response. Molecules 23: 3205.

Fousteri M, Mullenders LH. 2008. Transcription-coupled nucleotide excision repair in mammalian cells: molecular mechanisms and biological effects. Cell Res 18: 73–84.

Georges A, Gopaul D, Denby Wilkes C, Giordanengo Aiach N, Novikova E, Barrault MB, Alibert O, Soutourina J. 2019. Functional interplay between Mediator and RNA polymerase II in Rad2/XPG loading to the chromatin. Nucleic Acids Res 47: 8988–9004.

Hanawalt PC, Spivak G. 2008. Transcription-coupled DNA repair: two decades of progress and surprises. Nat Rev Mol Cell Biol 9: 958–970.

Holstege FC, Jennings EG, Wyrick JJ, Lee TI, Hengartner CJ, Green MR, Golub TR, Lander ES, Young RA. 1998. Dissecting the regulatory circuitry of a eukaryotic genome. Cell 95: 717–728.

Jeronimo C, Langelier MF, Bataille AR, Pascal JM, Pugh BF, Robert F. 2016. Tail and Kinase Modules Differently Regulate Core Mediator Recruitment and Function In Vivo. Molecular cell 64: 455–466.

Jeronimo C, Robert F. 2014. Kin28 regulates the transient association of Mediator with core promoters. Nature structural & molecular biology 21: 449–455.

Kamileri I, Karakasilioti I, Sideri A, Kosteas T, Tatarakis A, Talianidis I, Garinis GA. 2012. Defective transcription initiation causes postnatal growth failure in a mouse model of nucleotide excision repair (NER) progeria. Proc Natl Acad Sci U S A 109: 2995–3000.

Kaufman L, Rousseeuw PJ. 1990. Finding Groups in Data: An Introduction to Cluster Analysis. Hoboken, NJ: John Wiley & Sons, Inc.

Kikuchi Y, Umemura H, Nishitani S, Iida S, Fukasawa R, Hayashi H, Hirose Y, Tanaka A, Sugasawa K, Ohkuma Y. 2015. Human mediator MED17 subunit plays essential roles in gene regulation by associating with the transcription and DNA repair machineries. Genes Cells 20: 191–202.

Kokic G, Wagner FR, Chernev A, Urlaub H, Cramer P. 2021. Structural basis of human transcription-DNA repair coupling. Nature 598: 368–372.

Kornberg RD. 2005. Mediator and the mechanism of transcriptional activation. Trends Biochem Sci 30: 235–239.

Lake RJ, Boetefuer EL, Tsai PF, Jeong J, Choi I, Won KJ, Fan HY. 2014. The sequence-specific transcription factor c-Jun targets Cockayne syndrome protein B to regulate transcription and chromatin structure. PLoS Genet 10: e1004284.

Langmead B, Salzberg SL. 2012. Fast gapped-read alignment with Bowtie 2. Nat Methods 9: 357–359.

Le May N, Fradin D, Iltis I, Bougneres P, Egly JM. 2012. XPG and XPF endonucleases trigger chromatin looping and DNA demethylation for accurate expression of activated genes. Molecular cell 47: 622–632.

Le May N, Mota-Fernandes D, Velez-Cruz R, Iltis I, Biard D, Egly JM. 2010. NER factors are recruited to active promoters and facilitate chromatin modification for transcription in the absence of exogenous genotoxic attack. Molecular cell 38: 54–66.

Li H, Handsaker B, Wysoker A, Fennell T, Ruan J, Homer N, Marth G, Abecasis G, Durbin R, Genome Project Data Processing S. 2009. The Sequence Alignment/Map format and SAMtools. Bioinformatics 25: 2078–2079.

Linder T, Zhu X, Baraznenok V, Gustafsson CM. 2006. The classical srb4-138 mutant allele causes dissociation of yeast Mediator. Biochem Biophys Res Commun 349: 948–953.

Malabat C, Feuerbach F, Ma L, Saveanu C, Jacquier A. 2015. Quality control of transcription start site selection by nonsense-mediated-mRNA decay. Elife 4: e06722.

Malik S, Chaurasia P, Lahudkar S, Durairaj G, Shukla A, Bhaumik SR. 2010. Rad26p, a transcription-coupled repair factor, is recruited to the site of DNA lesion in an elongating RNA polymerase II-dependent manner in vivo. Nucleic Acids Res 38: 1461–1477.

Malik S, Chaurasia P, Lahudkar S, Uprety B, Bhaumik SR. 2012. Rad26p regulates the occupancy of histone H2A-H2B dimer at the active genes in vivo. Nucleic Acids Res 40: 3348–3363.

McNeil EM, Melton DW. 2012. DNA repair endonuclease ERCC1-XPF as a novel therapeutic target to overcome chemoresistance in cancer therapy. Nucleic Acids Res 40: 9990–10004.

Menck CF, Munford V. 2014. DNA repair diseases: What do they tell us about cancer and aging? Genet Mol Biol 37: 220–233.

Menoni H, Wienholz F, Theil AF, Janssens RC, Lans H, Campalans A, Radicella JP, Marteijn JA, Vermeulen W. 2018. The transcription-coupled DNA repair-initiating protein CSB promotes XRCC1 recruitment to oxidative DNA damage. Nucleic Acids Res 46: 7747–7756.

Mullenders L. 2015. DNA damage mediated transcription arrest: Step back to go forward. DNA repair 36: 28–35.

Newman JC, Bailey AD, Weiner AM. 2006. Cockayne syndrome group B protein (CSB) plays a general role in chromatin maintenance and remodeling. Proc Natl Acad Sci U S A 103: 9613–9618.

Nonet M, Scafe C, Sexton J R Y. 1987. Eucaryotic RNA polymerase conditional mutant that rapidly ceases mRNA synthesis. Mol Cell Biol 7: 1602–1611.

Pascucci B, Fragale A, Marabitti V, Leuzzi G, Calcagnile AS, Parlanti E, Franchitto A, Dogliotti E, D’Errico M. 2018. CSA and CSB play a role in the response to DNA breaks. Oncotarget 9: 11581–11591.

Paul E, Zhu Z, Landsman D, Morse R. 2015. Genome-wide association of mediator and RNA polymerase II in wild-type and mediator mutant yeats. Mol Cell Biol 35: 331–342.

Pelechano V, Wei W, Steinmetz LM. 2013. Extensive transcriptional heterogeneity revealed by isoform profiling. Nature 497: 127–131.

Petrenko N, Jin Y, Wong KH, Struhl K. 2016. Mediator Undergoes a Compositional Change during Transcriptional Activation. Molecular cell 64: 443–454.

Proietti-De-Santis L, Drane P, Egly JM. 2006. Cockayne syndrome B protein regulates the transcriptional program after UV irradiation. EMBO J 25: 1915–1923.

Ramirez F, Ryan DP, Gruning B, Bhardwaj V, Kilpert F, Richter AS, Heyne S, Dundar F, Manke T. 2016. deepTools2: a next generation web server for deep-sequencing data analysis. Nucleic Acids Res 44: W160–165.

Rockx DA, Mason R, van Hoffen A, Barton MC, Citterio E, Bregman DB, van Zeeland AA, Vrieling H, Mullenders LH. 2000. UV-induced inhibition of transcription involves repression of transcription initiation and phosphorylation of RNA polymerase II. Proc Natl Acad Sci U S A 97: 10503–10508.

Rousseeuw PJ. 1987. Silhouettes: A graphical aid to the interpretation and validation of cluster analysis. Journal of Computational and Applied Mathematics 20: 53–65.

Sabatella M, Theil AF, Ribeiro-Silva C, Slyskova J, Thijssen K, Voskamp C, Lans H, Vermeulen W. 2018. Repair protein persistence at DNA lesions characterizes XPF defect with Cockayne syndrome features. Nucleic Acids Res 46: 9563–9577.

Santos-Pereira JM, Aguilera A. 2015. R loops: new modulators of genome dynamics and function. Nat Rev Genet 16: 583–597.

Sarker AH, Tsutakawa SE, Kostek S, Ng C, Shin DS, Peris M, Campeau E, Tainer JA, Nogales E, Cooper PK. 2005. Recognition of RNA polymerase II and transcription bubbles by XPG, CSB, and TFIIH: insights for transcription-coupled repair and Cockayne Syndrome. Molecular cell 20: 187–198.

Schiano C, Casamassimi A, Rienzo M, de Nigris F, Sommese L, Napoli C. 2014. Involvement of Mediator complex in malignancy. Biochim Biophys Acta 1845: 66–83.

Selby CP, Sancar A. 1997a. Cockayne syndrome group B protein enhances elongation by RNA polymerase II. Proc Natl Acad Sci U S A 94: 11205–11209.

Selby CP, Sancar A. 1997b. Human transcription-repair coupling factor CSB/ERCC6 is a DNA-stimulated ATPase but is not a helicase and does not disrupt the ternary transcription complex of stalled RNA polymerase II. J Biol Chem 272: 1885–1890.

Sollier J, Stork CT, Garcia-Rubio ML, Paulsen RD, Aguilera A, Cimprich KA. 2014. Transcription-coupled nucleotide excision repair factors promote R-loop-induced genome instability. Molecular cell 56: 777–785.

Soutourina J. 2018. Transcription regulation by the Mediator complex. Nat Rev Mol Cell Biol 19: 262–274.

Spaeth JM, Kim NH, Boyer TG. 2011. Mediator and human disease. Semin Cell Dev Biol 22: 776–787.

Takagi Y, Kornberg RD. 2006. Mediator as a general transcription factor. Journal of Biological Chemistry 281: 80–89.

Teng Y, Yadav T, Duan M, Tan J, Xiang Y, Gao B, Xu J, Liang Z, Liu Y, Nakajima S et al. 2018. ROS-induced R loops trigger a transcription-coupled but BRCA1/2-independent homologous recombination pathway through CSB. Nat Commun 9: 4115.

Thompson CM, Young RA. 1995. General requirement for RNA polymerase II holoenzymes in vivo. Proc Natl Acad Sci U S A 92: 4587–4590.

van den Boom V, Citterio E, Hoogstraten D, Zotter A, Egly JM, van Cappellen WA, Hoeijmakers JH, Houtsmuller AB, Vermeulen W. 2004. DNA damage stabilizes interaction of CSB with the transcription elongation machinery. J Cell Biol 166: 27–36.

van Gool AJ, Verhage R, Swagemakers SM, van de Putte P, Brouwer J, Troelstra C, Bootsma D, Hoeijmakers JH. 1994. RAD26, the functional S. cerevisiae homolog of the Cockayne syndrome B gene ERCC6. EMBO J 13: 5361–5369.

van Vuuren AJ, Appeldoorn E, Odijk H, Yasui A, Jaspers NG, Bootsma D, Hoeijmakers JH. 1993. Evidence for a repair enzyme complex involving ERCC1 and complementing activities of ERCC4, ERCC11 and xeroderma pigmentosum group F. EMBO J 12: 3693–3701.

Vannier JB, Depeiges A, White C, Gallego ME. 2009. ERCC1/XPF protects short telomeres from homologous recombination in Arabidopsis thaliana. PLoS Genet 5: e1000380.

Vermeij WP, Dolle ME, Reiling E, Jaarsma D, Payan-Gomez C, Bombardieri CR, Wu H, Roks AJ, Botter SM, van der Eerden BC et al. 2016. Restricted diet delays accelerated ageing and genomic stress in DNA-repair-deficient mice. Nature 537: 427–431.

Wahba L, Costantino L, Tan FJ, Zimmer A, Koshland D. 2016. S1-DRIP-seq identifies high expression and polyA tracts as major contributors to R-loop formation. Genes & Development 30: 1327–1338.

Wang W, Xu J, Chong J, Wang D. 2018. Structural basis of DNA lesion recognition for eukaryotic transcription-coupled nucleotide excision repair. DNA repair 71: 43–55.

Wang Y, Chakravarty P, Ranes M, Kelly G, Brooks PJ, Neilan E, Stewart A, Schiavo G, Svejstrup JQ. 2014. Dysregulation of gene expression as a cause of Cockayne syndrome neurological disease. Proc Natl Acad Sci U S A 111: 14454–14459.

Wei L, Nakajima S, Bohm S, Bernstein KA, Shen Z, Tsang M, Levine AS, Lan L. 2015. DNA damage during the G0/G1 phase triggers RNA-templated, Cockayne syndrome B-dependent homologous recombination. Proc Natl Acad Sci U S A 112: E3495–3504.

Wickham H. 2009. ggplot2: Elegant Graphics for Data Analysis. Springer-Verlag New York.

Wilkinson L. 2006. Revising the Pareto Chart. The American Statistician 60: 332–334.

Wong KH, Jin Y, Struhl K. 2014. TFIIH Phosphorylation of the Pol II CTD Stimulates Mediator Dissociation from the Preinitiation Complex and Promoter Escape. Molecular cell 54: 601–612.

Xu J, Lahiri I, Wang W, Wier A, Cianfrocco MA, Chong J, Hare AA, Dervan PB, DiMaio F, Leschziner AE et al. 2017. Structural basis for the initiation of eukaryotic transcription-coupled DNA repair. Nature 551: 653–657.

Xu J, Wang W, Xu L, Chen J, Chong J, Oh J, Leschziner A, Fu X-D, Wang D. 2020. Cockayne syndrome B protein acts as an ATP-dependent processivity factor that helps RNA polymerase II overcome barriers. PNAS.

Zaros C, Thuriaux P. 2005. Rpc25, a conserved RNA polymerase III subunit, is critical for transcription initiation. Molecular Microbiology 55: 104–114.

Zhang Y, Liu T, Meyer CA, Eeckhoute J, Johnson DS, Bernstein BE, Nusbaum C, Myers RM, Brown M, Li W et al. 2008. Model-based analysis of ChIP-Seq (MACS). Genome Biol 9: R137.

Zhu XD, Niedernhofer L, Kuster B, Mann M, Hoeijmakers JH, de Lange T. 2003. ERCC1/XPF removes the 3’ overhang from uncapped telomeres and represses formation of telomeric DNA-containing double minute chromosomes. Molecular cell 12: 1489–1498.

