## Supplementary Data for "Genome-wide distribution of Rad26 and Rad1-Rad10 reveals their relationship with Mediator and RNA polymerase II"

**Supplementary Table S1. Yeast strains used in the study.**

| Collection # | Strain Name | Genotype |
|---|---|---|
| **Y0097** | YPH499 WT | *MATa ura3-52 his3-Δ200 ade2-101uaa trp1-Δ63 lys2-801uag leu2-Δ1* |
| **Y6611** | BY4741 WT | *MATa his3-Δ1 leu2Δ-0 met15Δ-0 ura3Δ-0* |
| **Y6612** | BY4742 WT | *MATα his3Δ-1 leu2Δ-0 lys2Δ-0 ura3Δ-0* |
| **Y6244** | YPH499 rad26Δ | *MATa ura3-52 his3-Δ200 ade2-101uaa trp1-Δ63 lys2-801uag leu2-Δ1 rad26::KanMX6* |
| **Y7726** | BY4742 rad26Δ | *MATα his3Δ-1 leu2Δ-0 lys2Δ-0 ura3Δ-0 rad26::KanMX6* |
| **Y6245** | YPH499 rad7Δ | *MATa ura3-52 his3-Δ200 ade2-101uaa trp1-Δ63 lys2-801uag leu2-Δ1 rad7::HIS3* |
| **Y7116** | BY4742 rad7Δ | *MATα his3Δ-1 leu2Δ-0 lys2Δ-0 ura3Δ-0 rad7::KanMX6* |
| **Y6247** | YPH499 rad26Δ rad7Δ | *MATa ura3-52 his3-Δ200 ade2-101uaa trp1-Δ63 lys2-801uag leu2-Δ1 rad26::KanMX6 rad7::HIS3* |
| **Y7727** | BY4742 rad26Δ rad7Δ | *MATα his3Δ-1 leu2Δ-0 lys2Δ-0 ura3Δ-0 rad26::KanMX6 rad7::HIS3* |
| **Y7314** | YPH499 rad10Δ | *MATa ura3-52 his3-Δ200 ade2-1 trp1-Δ63 lys2-801uag leu2-Δ rad10::hph* |
| **Y6191** | BY4741 rad10Δ | *MATa his3-Δ leu2Δ-0 met15Δ-0 ura3Δ-0 rad10::KanMX4* |
| **Y7313** | YPH499 rad1Δ | *MATa ade2-1 lys2-801 ura3-52 trp1-Δ63 his3-Δ200 leu2-Δ rad1::hph* |
| **Y6187** | BY4741 rad1Δ | *MATa his3-Δ1 leu2Δ-0 met15Δ-0 ura3Δ-0 rad1::KanMX4* |
| **Y4582** | dst1Δ | *MATa lys2-801 trp1-Δ63 his3-Δ200 leu2-Δ1 dst1::KanMX4* |
| **Y6200** | Rad1-HA | *MATa ura3-52 his3-Δ200 ade2-101uaa trp1-Δ63 lys2-801uag leu2-Δ1 RAD1::3HA::HIS3* |
| **Y6201** | HA-Rad1 | *MATa ura3-52 his3-Δ200 ade2-101uaa trp1-Δ63 lys2-801uag leu2-Δ1 HIS3::pADH1::3HA::RAD1* |
| **Y6204** | Rad10-HA | *MATa ura3-52 his3-Δ200 ade2-101uaa trp1-Δ63 lys2-801uag leu2-Δ1 RAD10::3HA::HIS3* |
| **Y7064** | Rad10-Flag | *MATa ade2-1 lys2-801 ura3-52 trp1-Δ63 his3-Δ200 leu2Δ RAD10::3Flag::KanMX4* |
| **Y6572** | HA-Rad1 Med17-Myc | *MATa ura3-52 his3-Δ200 ade2-101uaa trp1-Δ63 lys2- Δ1 leu2-Δ1 HIS3::pADH1::3HA::RAD1 MED17::13MYC::TRP1* |
| **Y7061** | Rad10-Flag Med17-Myc | *MATα ura3-52 his3-Δ200 ade2-101uaa trp1-Δ63 lys2- 801uag leu2-Δ1 RAD10::3Flag::KanMX4 MED17::13MYC::TRP1* |
| **Y7050** | HA-Rad26 | *MATa ade2-1 lys2-801 ura3-52 trp1-Δ63 his3-Δ200 leu2Δ HIS3::HA::RAD26* |
| **Y7194** | HA-Rad26 Med17-Myc | *MATa ade2-1 lys2-801 ura3-52 trp1-Δ63 his3-Δ200 leu2Δ HIS3::HA::RAD26 MED17::13MYC::TRP1* |
| **Y7629** | rpb1-1 Rad10-HA | *MATα ura3-52 leu2-Δ1 his3-Δ200 trp1-Δ63 rpb1-1(rpb1-G1437D) RAD10::3HA::HIS* |
| **Y7628** | rpb1-1 Rad1-HA | *MATα ura3-52 leu2-Δ1 his3-Δ200 trp1-Δ63 rpb1-1(rpb1-G1437D) RAD1::3HA::HIS3* |
| **Y7117** | Rad26-HA rad7Δ | *MATa ura3-52 his3-Δ200 ade2-101uaa trp1-Δ63 lys2-801uag leu2-Δ1 RAD26::3HA::KanMX6 rad7::HIS* |
| **Y7118** | HA-Rad26 rad7Δ | *MATa ade2-1 lys2-801 ura3-52 trp1-Δ63 his3-Δ200 leu2Δ HIS3::HA::RAD26 rad7::KAN* |
| **Y6202** | Rad14-HA | *MATa ura3-52 his3-Δ200 ade2-101uaa trp1-Δ63 lys2-801uag leu2-Δ1 RAD14::3HA::HIS3* |
| **Y6203** | HA-Rad14 | *MATa ura3-52 his3-Δ200 ade2-101uaa trp1-Δ63 lys2-801uag leu2-Δ1 HIS3::pADH1::3HA::RAD14* |

| | | |
|---|---|---|
| **Y6578** | Rad4-HA | *MATa ade2-1 lys2-801 ura3-52 trp1-Δ63 his3-Δ200 leu2Δ RAD4::HA::HIS3* |
| **Y6579** | HA-Rad4 | *MATa ade2-1 lys2-801 ura3-52 trp1-Δ63 his3-Δ200 leu2Δ HIS3::HA::RAD4* |
| **Y6573** | Rad14-HA<br>Med17-Myc | *MATa ura3-52 his3-Δ200 ade2-101uaa trp1-Δ63 lys2- 801uag leu2-Δ1 RAD14::3HA::HIS3 MED17::13MYC::TRP1* |
| **Y6574** | HA-Rad14<br>Med17-Myc | *MATa ura3-52 his3-Δ200 ade2-101uaa trp1-Δ63 lys2- 801uag leu2-Δ1 HIS3::pADH1::3HA::RAD14 MED17::13MYC::TRP1* |
| **Y6604** | Rad4-HA<br>Med17-Myc | *MATa ade2-1 lys2- 801 ura3-52 trp1-Δ63 his3-Δ200 leu2-Δ l RAD4::3HA::HIS3 MED17::13MYC::TRP1* |
| **Y6605** | HA-Rad4<br>Med17-Myc | *MATa ade2-1 lys2-801 ura3-52 trp1-Δ63 his3-Δ200 leu2-Δ HIS3::pADH1::3HA::RAD4 MED17::13MYC::TRP1* |
| **Y6300** | YPH500<br>Med17-Myc | *MATα ura3-52 his3-Δ200 ade2-101uaa trp1-Δ63 lys2-801uag leu2-Δ1 MED17::13MYC::TRP1* |
| **Y7751** | YPH499<br>Med17-Myc | *MATa ura3-52 his3-Δ200 ade2-101uaa lys2-801uag leu2-Δ1 MED17::13Myc::TRP1* |
| **Y7641** | Rad10-HA<br>rpc25-S100P | *MATa ura3-52 trp1-Δ63 his3- 200 leu2- Δ1 ade2-101uaa lys2-801 ade2-1 rpc25-S100P RAD10::3HA::HIS3* |
| **Y7639** | Rad10-Flag<br>rad2Δ | *MATα ura3-52 his3- Δ200 ade2-101uaa trp1-Δ63 lys2-801uag leu2-Δ1 rad2::KanMX6 RAD10::3Flag::hph* |
| **Y7734** | Rad10-HA<br>Med5-Myc<br>MED17-WT | *MATa ura3-52 his3-Δ200 ade2-101uaa trp1-Δ63 lys2-801uag leu2-Δ1 Med17::kan::ADE2 MED5::13Myc::KanMX6 RAD10::3HA::HIS3 CEN MED17 LEU2* |
| **Y7735** | Rad10-HA<br>Med5-Myc<br>med17-138 | *MATa ura3-52 his3-Δ200 ade2-101uaa trp1-Δ63 lys2-801uag leu2-Δ1 med17::kan::ADE2 MED5::13Myc::KanMX6 RAD10::3HA::HIS3 CEN med17-138 LEU2* |
| **Y7730** | HA-Rad26<br>Med5-Myc<br>MED17-WT | *MATa ura3-52 his3-Δ200 ade2-101uaa trp1-Δ63 lys2-801uag leu2-Δ1 med17::kan::ADE2 MED5::13Myc::KanMX6 HIS ::3HA::RAD26 CEN MED17 LEU2* |
| **Y7731** | HA-Rad26<br>Med5-Myc<br>med17-138 | *MATa ura3-52 his3-Δ200 ade2-101uaa trp1-Δ63 lys2-801uag leu2-Δ1 med17::kan::ADE2 MED5::13Myc::KanMX6 HIS ::3HA::RAD26 CEN med17-138 LEU2* |
| **Y7666** | Rad10-HA<br>Med17-Myc<br>KIN28-WT | *MATα ade2-1 lys2-801uag ura3-52 trp1-Δ63 his3-Δ200 leu2-Δ RAD10::3HA::HIS3 MED17::13Myc::TRP1 kin28::KanMX6 CEN KIN28 LEU2* |
| **Y7667** | Rad10-HA<br>Med17-Myc<br>kin28-ts16 | *MATα ade2-1 lys2-801uag ura3-52 trp1-Δ63 his3-Δ200 leu2-Δ RAD10::3HA::HIS3 MED17::13Myc::TRP1 kin28::KanMX6 CEN kin28-ts16 LEU2* |
| **Y7668** | HA-Rad26<br>Med17-Myc<br>KIN28-WT | *MATα ade2-1 lys2-801uag ura3-52 trp1-Δ63 his3-Δ200 leu2-Δ HIS3::3HA::RAD26 MED17::13Myc::TRP1 kin28::KanMX6 CEN KIN28 LEU2* |
| **Y7669** | HA-Rad26<br>Med17-Myc<br>kin28-ts | *MATα ade2-1 lys2-801uag ura3-52 trp1-Δ63 his3-Δ200 leu2-Δ HIS3::3HA::RAD26 MED17::13Myc::TRP1 kin28::KanMX6 CEN kin28-ts16 LEU2* |
| **Y6960** | Rad1-HA<br>Med17-Myc<br>KIN28-WT | *MATa ura3-52 his3-Δ200 ade2-101uaa trp1-Δ63 lys2-801uag leu2-Δ1 RAD1::3HA::HIS3 MED17::13Myc::TRP1 kin28::KanMX6 CEN KIN28 LEU2* |
| **Y6961** | Rad1-HA<br>Med17-Myc<br>kin28-ts16 | *MATa ura3-52 his3-Δ200 ade2-101uaa trp1-Δ63 lys2-801uag leu2-Δ1 RAD1::3HA::HIS3 MED17::13Myc::TRP1 kin28::KanMX6 CEN kin28ts16 LEU2* |
| **Y7196** | Rad2-HA<br>Med17-Myc<br>rad10Δ | *MATα ade2-1 lys2-801 ura3-52 trp1-Δ63 his3-Δ200 leu2-Δ HIS3::pADH1::3HA::RAD2 rad10::KanMX6 MED17::13Myc::TRP1* |
| **Y7195** | Rad2-HA<br>Med17-Myc<br>rad1Δ | *MATα ade2-1 lys2-801 ura3-52 trp1-Δ63 his3-Δ200 leu2-Δ HIS3::pADH1::3HA::RAD2 rad1::KanMX6 MED17::13Myc::TRP1* |
| **Y7752** | Med17-Myc<br>rad26Δ | *MATa ura3-52 his3-Δ200 ade2-101uaa trp1-Δ63 lys2-801uag leu2-Δ1 rad26::KanMX6 MED17::13Myc::TRP1* |

**Supplementary Table S2. qPCR primers used in the study.**

| Name | Forward | Reverse |
|---|---|---|
| *ADH1-U* | ATAGGCGCATGCAACTTCTT | CATCAGCTCTGGAACAACGA |
| *CLN1-U* | ATTCCCTTGTTCGCAACACT | CGCGGGGTTGTAGTAGGTAA |
| *LEU1-U* | CGTCATAATAGTTCCTGCCCAGCT | TGTATCTTGAGTGTCTGTATGGGCG |
| *PCL1-U* | GGCTCTAGTGTTCTGCGAATG | ACATTTCGAGTCGCTTTTGC |
| *PDR5-U* | AAGACTGCCCCTCTCTTTCC | AAGACTGCCCCTCTCTTTCC |
| *PMA1-U* | AACAAACCCGGTCTCGAAG | GAAGTGCCGCATTAGGAAAT |
| *PYK1-U* | CACCGTCACAAAGTGTT | TGGGAAGGAAAGGAAATCAC |
| *SSF2-U* | GCGAATGGCTTGTGAAACAAACA | TGCAAGTGCTTAGGTCTGTTTGG |
| *OLE1-U* | TGACAGGCAGAGGTAATAACGG | AAGACTCGTAGAAGCACACCTG |
| *PCK1-U* | CAATTCTGACCAGAGCACTTGG | CAGATCTCCATTCACCCGTCAG |
| *ADH1-P* | TTCCTTCATTCACGCACACT | AGGGAACGAGAACAATGACG |
| *PMA1-P* | GATGGTGGGTACCGCTTATG | TTGGTGTTATAGGAAAGAAAGAGAAAA |
| *PYK1-P* | CCTTTCCTTCCCATATGATGC | ACTTTGAAAGGGGACCATGA |
| *SGV1-P* | CGCGTCGAACGGAAATACACCC | CTGCTTGACCACGAATGTACTGC |
| *ADH1-O* | GGCTGGAAGATCGGTGACTA | TCAGCGGTAGCGTATTGTTG |
| *BRF2-O* | GGTGGAAAAGTCTAAGTTAAGAGCAG | CGTCCTCATTCTCGGAAACTTC |
| *GAL1-O* | AAAGAAACTTGCACCGGAAA | GGCCCATATTCGCTTTAACA |
| *PDR5-O* | CTCCAGGCTATGACCCAAAA | GTTCTTCCAAGCGCAACCTA |
| *PMA1-O* | TCTCCAAAGCCCGTTAAATG | CGTTCATAGCACCGAAGTT |
| *PYK1-O* | ATGGTTGCCAGAGGTGACTT | TCTGGTTGGTCTTGGGTTGT |
| *TPI1-O* | TAACGTCGTTGTCGCTTACG | CCAACTTGGAAGCCAAGAAC |
| *RAD2-O* | TGAGTCTTCTAACGCGACGA | TAAGGCAACCTTGCAGCTTT |
| *IGV* | TTCATTTGCAATTTGCAGTTCA | CAGCCAGGAAGAATCTCACAA |
| *TEL1R* | TCAACTACCCTCCCTCTCCAT | GTGGAGTGGGGGAATGAGAC |
| *TEL1R-XC1* | CATCTATCCCCTGCCCAATA | ACGTGGTAGATGGGGATTGT |
| *RPR1* | ATGGTACGCTGTGGTGCTC | CCATAGGTGGGGATCCTTCT |
| *tDNAMet* | GCTTCAGTAGCTCAGTAGGAA | TGCTCCAGGGGAGGTTC |
| *tDNAPhe* | GACGCTTGGACCATTTATAAAGCAC | CCATAAGAGAAGGAGCAGTCAAGTTCA |
| *SCR1* | ACCGCTGTTAGGGGAGTTTT | CCAAATTAAACCGCCGAAG |

**Supplementary Table S3. Number of mapped and unique reads, and normalization coefficients in ChIP-seq experiments.**

| Name | Unique mapped reads | Average Insertion Size | Normalization Factor |
|---|---|---|---|
| Input_Kin28WT | 27621977 | 321 | 1 |
| Input_Kin28TS | 22653444 | 309 | 1 |
| Rad26_Kin28WT | 20512333 | 250 | 1 |
| Rad26_Kin28TS | 15914202 | 218 | 1.6515 |
| Med17_Kin28WT | 13479060 | 191 | 1 |
| Med17_Kin28TS | 9728462 | 205 | 0.9861 |
| Pol2_Kin28WT | 20472442 | 249 | 1 |
| Pol2_Kin28TS | 20810040 | 199 | 3.9843 |
| Rad1_Kin28WT | 13949359 | 216 | 1 |
| Rad1_Kin28TS | 8367165 | 175 | 0.8090 |
| Rad10_Kin28WT | 18118057 | 198 | 1 |
| Rad10_Kin28TS | 17453391 | 186 | 0.7715 |

#### Supplementary Figure Legends

**Supplementary Figure S1. UV-sensitivity of *rad26* deletion and Rad26, Rad1 and Rad10-tagged strains.**

Wild-type and isogenic deletion or tagged strains for Rad26 (GGR-proficient or deficient *rad7*Δ contexts, **A**) or tagged strains for Rad1 and Rad10 (**B**) were serially diluted, spotted on rich-medium YPD agar plates, treated or not with 20J/$m^2$ of UV (UV Stratalinker 1800) and incubated for 3 days at 30°C. The UV sensitivity of Rad26 strains was needed to be tested in a GGR-deficient context, since *rad26* deletion alone did not lead to UV sensitivity in yeast. Rad1-Rad10 are common for both NER subpathways and the UV sensitivity of Rad1-Rad10 tagged strains can thus be tested in a wild-type context.

**Supplementary Figure S2. ChIP analysis of Rad26, Rad1 and Rad10 occupancy on selected regions.**

Quantitative ChIP analysis of Rad26, Rad1 and Rad10 occupancy was performed with α-HA antibody against HA-tagged Rad26 (**A**), Rad1 (**B**) or Rad10 (**C**). Yeast strains carrying N-terminal HA-tagged versions of Rad26 (**A**), Rad1 (**B**, right panel) or Rad10 (**C**, right panel), or carrying C-terminal HA-tagged versions of Rad1 (**B**, left panel) or Rad10 (**C**, left panel) were grown in YPD medium at 30°C. Non-tagged (NT) strain was also analyzed (**A, D**). Immunoprecipitated fragments from ChIP experiments were amplified with primers corresponding to selected class II gene promoters (P), UAS (U) or ORFs (O), or class III *RPR1* gene or telomere *TEL1R*. A *GAL1* ORF and a non-transcribed region on chromosome V (IGV) were used as negative controls. Mean values and standard deviation (indicated by error bars) of three independent experiments are shown. Background is represented as a dotted line.

**Supplementary Figure S3. Genome-wide analysis of Rad26, Rad1 and Rad10.**

(**A**) Pie charts showing the distribution of Rad26, Rad1 and Rad10 enrichment peaks within annotated genomic features including Pol II transcribed regions of protein-coding genes (mRNAs), intergenic regions, telomeres, centromeres, Pol III-transcribed tRNA genes. Proportion of these regions within the yeast genome is also shown.

(**B**) Heatmaps of input, Mediator (Med17), Pol II, Rad26, Rad1 and Rad10 ChIP-seq profiles on yeast centromeres (scaled windows for 500 bp before centromere, between start and end, and 500 bp after centromere), sorted by decreasing Rad10 occupancy. Average tag density (metagene) profiles in RPM (reads per million) are shown in upper panels. The centromere numbers are indicated on the left.

(**C**) Rad10 ChIP-seq density versus Rad1 ChIP-seq density on Pol III-transcribed tRNA genes. Each point on the plot corresponds to one gene. A linear regression (dotted line) and an $R^2$ linear regression coefficient are indicated. The dashed line corresponds to y = x.

**Supplementary Figure S4. Mediator and Pol II occupancy in *rad26*Δ strain and growth phenotypes of *rad26*Δ in GGR-proficient and deficient (*rad7*Δ) context.**

(**A, B**) Quantitative ChIP analysis of Mediator (Med17-Myc) and Pol II occupancy was performed with α-Myc antibody against Med17-Myc (**A**) or α-Rpb1 Pol II antibody (**B**) in WT and *rad26* deletion strains. Immunoprecipitated fragments from ChIP experiments were amplified with primers corresponding to selected class II gene UAS (U) or ORFs (O). A *GAL1* ORF and a non-transcribed region on chromosome V (IGV) were used as negative controls. Mean values and standard deviation (indicated by error bars) of three independent experiments are shown.

(**C**) Growth phenotypes of *rad26*Δ in GGR-proficient and deficient context.

Wild-type and isogenic *rad26* deletion strains in GGR-proficient or *rad7* (GGR-deficient) BY4741 and YPH499 contexts were serially diluted, spotted on indicated agar plates, and incubated for 3 days at 30°C. UV-sensitivity assays are shown in **Supplementary Figure S1**. Rich media supplemented with 2 or 0.05% glucose (YPD and YPD 0.05%, respectively), 2% galactose (YPGal), minimal SC medium containing indicated concentrations of mycophenolic acid (MPA), were used. *dst1* (TFIIS gene) deletion strain was added for MPA-sensitivity control.

**Supplementary Figure S5. Effect of *rpb1-1* mutation on Rad10 occupancy, growth phenotypes of *rad1* and *rad10* deletion strains and ChIP analysis of Pol II and Mediator in these deletion contexts.**

(**A, B**) Effect of *rpb1-1* Pol II mutation on Rad1 and Pol II occupancies on selected regions. Quantitative ChIP assays were performed using α-Rpb1 antibody (Pol II) (**A**), and α-HA antibody against Rad1-HA (**B**). Cells were grown in selective SD medium complemented with amino acids at 25°C and then shifted for 90 min at 37°C. *GAL1-O* amplicon was used as a negative control. Quantities were reported to qPCR performed on Input DNA and are expressed as a percentage. The indicated value is the mean of three biological replicates, and error bars represent the standard deviation.

(**C**) Growth phenotypes of *rad1*Δ and *rad10*Δ.

Wild-type and isogenic *rad1* and *rad10* deletion strains in BY4741 and YPH499 contexts were serially diluted, spotted on indicated agar plates, and incubated for 3 days at 30°C. For UV-sensitivity assays, cells were treated or not with 5J/m$^2$ of UV (UV Stratalinker 1800). Rich media supplemented with 2 or 0.05% glucose (YPD and YPD 0.05%, respectively), 2% galactose (YPGal), 1% ethanol (YPE 1%), 2% galactose (YPGalactose) or 2% glycerol (YPGlycerol), minimal SC medium containing indicated concentrations of mycophenolic acid

(MPA), were used. The only observed phenotype of *rad1* and *rad10* deletion strains (UV sensitivity) is highlighted in red.

(**D-G**) Mediator and Pol II ChIP analysis in *rad1* and *rad10* deletion strains on selected regions. Quantitative ChIP assays were performed using α-Rpb1 antibody (Pol II) (**D, F**), and α-Myc antibody against Med17-Myc (**E, G**). Wild-type and *rad1* (**D, E**) or *rad10* (**F, G**) deletion strains were grown in rich YPD medium at 30°C. A non-transcribed region on chromosome V (IGV) amplicon was used as a negative control. Quantities were reported to qPCR performed on Input DNA and are expressed as a percentage. The indicated value is the mean of three biological replicates, and error bars represent the standard deviation.

**Supplementary Figure S6. No coimmunoprecipitation between Mediator and Rad4 and Rad14.**

(**A**) Mediator was immunoprecipitated from crude yeast extracts via Med17-Myc subunit with α-Myc antibody (IP) and Western blotting with α-HA antibody detected Rad4-HA (CoIP).

(**B**) HA-Rad14 was immunoprecipitated with α-HA antibody from crude yeast extracts and analyzed by Western blotting with α-Myc antibody against Med17-Myc Mediator subunit. No coimmunoprecipitation with Med17 Mediator subunit was detected (CoIP) (left panels). IgG indicates a control immunoprecipitation with IgG magnetic beads only. Inputs are shown on right panels.

**Supplementary Figure S7. Effect of *med17-138* mutation on Rad10 and Rad26 occupancies on selected regions.**

Quantitative ChIP assays were performed using α-HA antibody against Rad10-HA (**A**) or HA-Rad26 (**B**). Cells were grown in selective SD medium complemented with amino acids at 25°C and then shifted for 45 min at 37°C. *GAL1-O* and *IGV* (non-transcribed region on chromosome

V) amplicons were used as negative controls. Quantities were reported to qPCR performed on Input DNA and are expressed as a percentage. The indicated value is the mean of three biological replicates, and error bars represent the standard deviation.

**Supplementary Figure S8. Rad26, Rad1, Rad10 ChIP analysis in comparison with Pol II and Mediator (Med17-Myc) in *kin28* context.**

Quantitative ChIP analysis of Mediator (Med17-Myc), Pol II, HA-Rad26, Rad10-HA and Rad1-HA occupancy was performed with α-Myc antibody against Med17-Myc (**A**), α-Rpb1 Pol II antibody (**B**) or α-HA antibody against HA-Rad26 (**C**), Rad10-HA (**D**) and Rad1-HA (**E**) in WT and *kin28-ts* strains. Cells were grown in selective SD medium complemented with amino acids at 25°C and then shifted for 75 min at 37°C. Immunoprecipitated fragments from ChIP experiments were amplified with primers corresponding to selected class II gene UAS (U), promoters (P) or ORFs (O). A *GAL1* ORF and a non-transcribed region on chromosome V (IGV) were used as negative controls. Mean values and standard deviation (indicated by error bars) of three independent experiments are shown.

**Supplementary Figure S9. Metagene analysis of genome-wide Mediator, Pol II, Rad26, Rad1 and Rad10 occupancy on Mediator enrichment peaks in WT and *kin28-ts* strains.**

Average tag density in WT strains is indicated as a full line, whereas average tag density in *kin28-ts* strains is indicated as a dashed line.

(**A, C**) Average tag density in Med17 Mediator, Pol II, Rad26, Rad1 and Rad10 ChIP and input, around Med17 Mediator enrichment peaks (-500 bp to +500 bp) determined in WT (**A**) or *kin28-ts* mutant (**C**).

(**B, D**) Med17 Mediator, Pol II, Rad26, Rad1 and Rad10 ChIP occupancy ratios between *kin28-ts* mutant and WT, around Med17 Mediator enrichment peaks (-500 bp to +500 bp) determined in WT (**A**) or *kin28-ts* mutant (**C**).

**Supplementary Figure S10. Metagene analysis of genome-wide Mediator, Pol II, Rad26, Rad1 and Rad10 occupancy on transcribed regions in WT and *kin28-ts* strains.**

(**A**) Heatmaps of Mediator (Med17), Pol II, Rad26, Rad1 and Rad10 ChIP-seq occupancy in WT and *kin28-ts* mutant on 10% Pol II-most enriched regions (scaled windows for 500 bp before TSS, between TSS and TES, and 500 bp after TES), sorted by decreasing Pol II occupancy.

(**B**) Med17 Mediator, Pol II, Rad26, Rad1 and Rad10 ChIP occupancy ratios between *kin28-ts* mutant and WT on 10% Pol II-most enriched regions (scaled windows for 500 bp before TSS, between TSS and TES, and 500 bp after TES).

**Supplementary Figure S11. PCA and clustering analysis of Mediator, Rad1, Rad10, Rad26 and Pol II ChIP-seq data on intergenic regions.**

(**A**) Pareto chart of the distribution of sample variance as a function of number of principle components (PCs) in WT (left panel) or *kin28-ts* (right panel) contexts. The histogram represents the explained variance in percentage (y-axis) for each PC (x-axis) and the line corresponds to the total variance explained by 1 to 4 PCs.

(**B**) Norms of each variable Med17 (Mediator), Pol II, Rad1, Rad10 and Rad26 in the orthonormal referential of 3PCs for WT and *kin28-ts* strains.

(**C**) Contribution of each variable to each PC (PC1, 2 and 3) in WT and *kin28-ts* strains. The values for main contributors are indicated in bold.

(**D** Silhouette values of k-list hierarchical clustering for WT (left panel) and *kin28-ts* (right panel) strains. The value is maximal when the data sets are separated in two independent clusters.

**Supplementary Figure S12. Projections of Mediator, Rad1, Rad10, Rad26 and Pol II ChIP-seq data on 2 PCA plane.**

Covariance of the variables Mediator (Med17), Pol II, Rad1, Rad10 and Rad26 for ChIP-seq data in intergenic regions were projected on 2 PCA axes (PC1 versus 2, PC2 versus 3, PC1 versus 3) in WT (**A**) or *kin28-ts* (**B**). Cluster 1 is shown in red and cluster 2 in blue.

**Supplementary Figure S13. Venn diagram for clusters in WT and *kin28-ts*.**

Venn diagram was generated for overlapping between clusters 1 and 2 in WT and *kin28-ts* mutant. The number of regions in each intersection is indicated.

**Supplementary Figure S1**

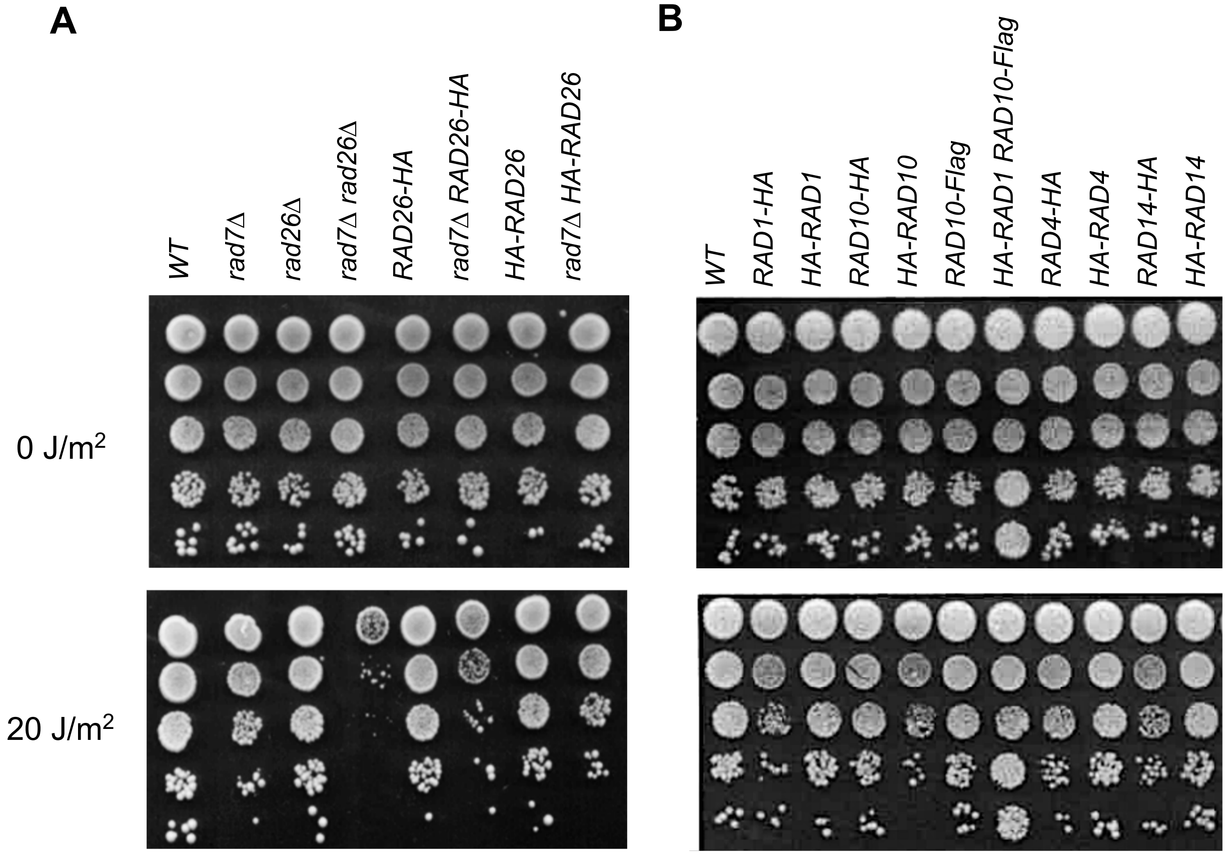

### Supplementary Figure S2

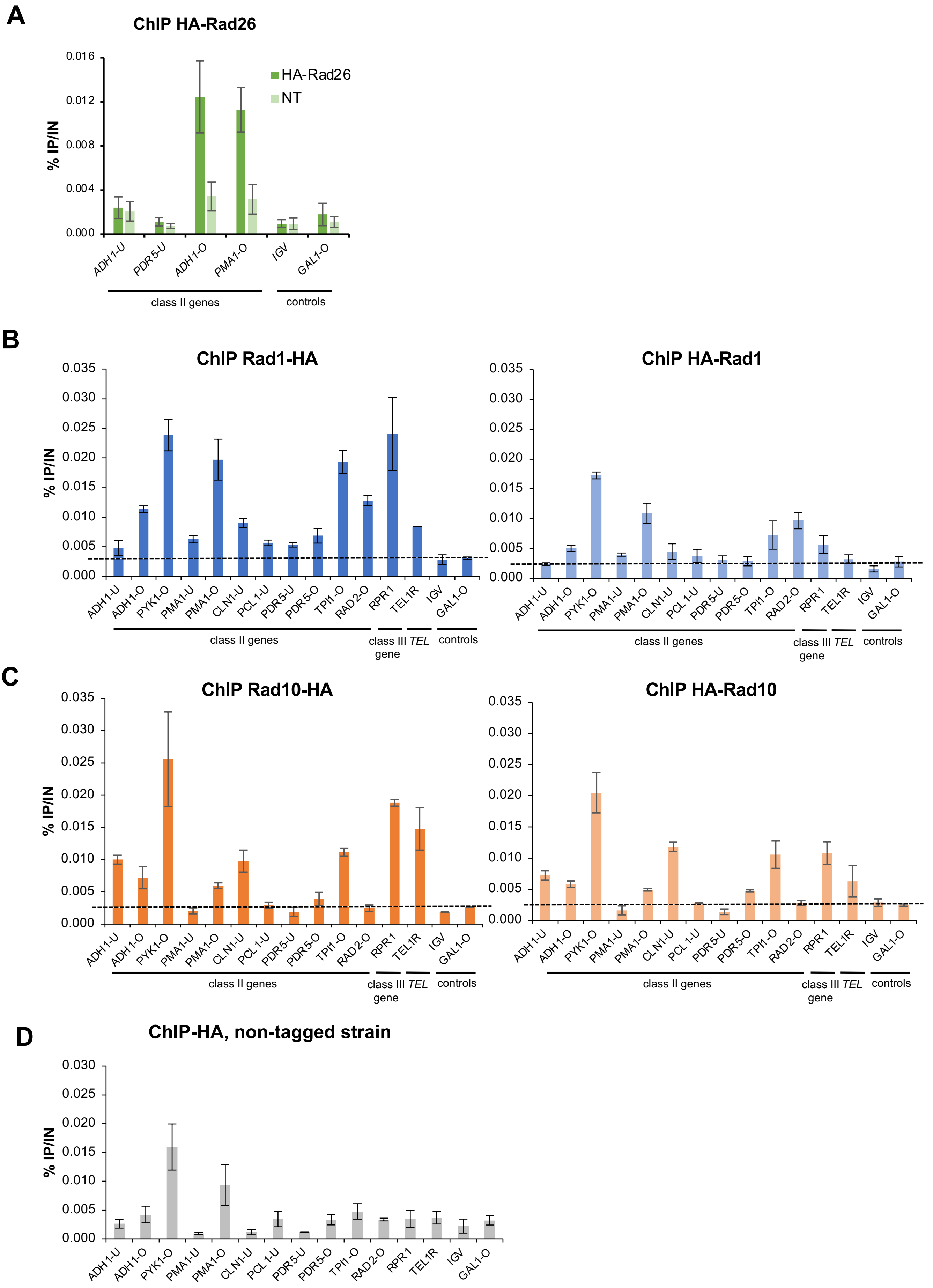

### Supplementary Figure S3

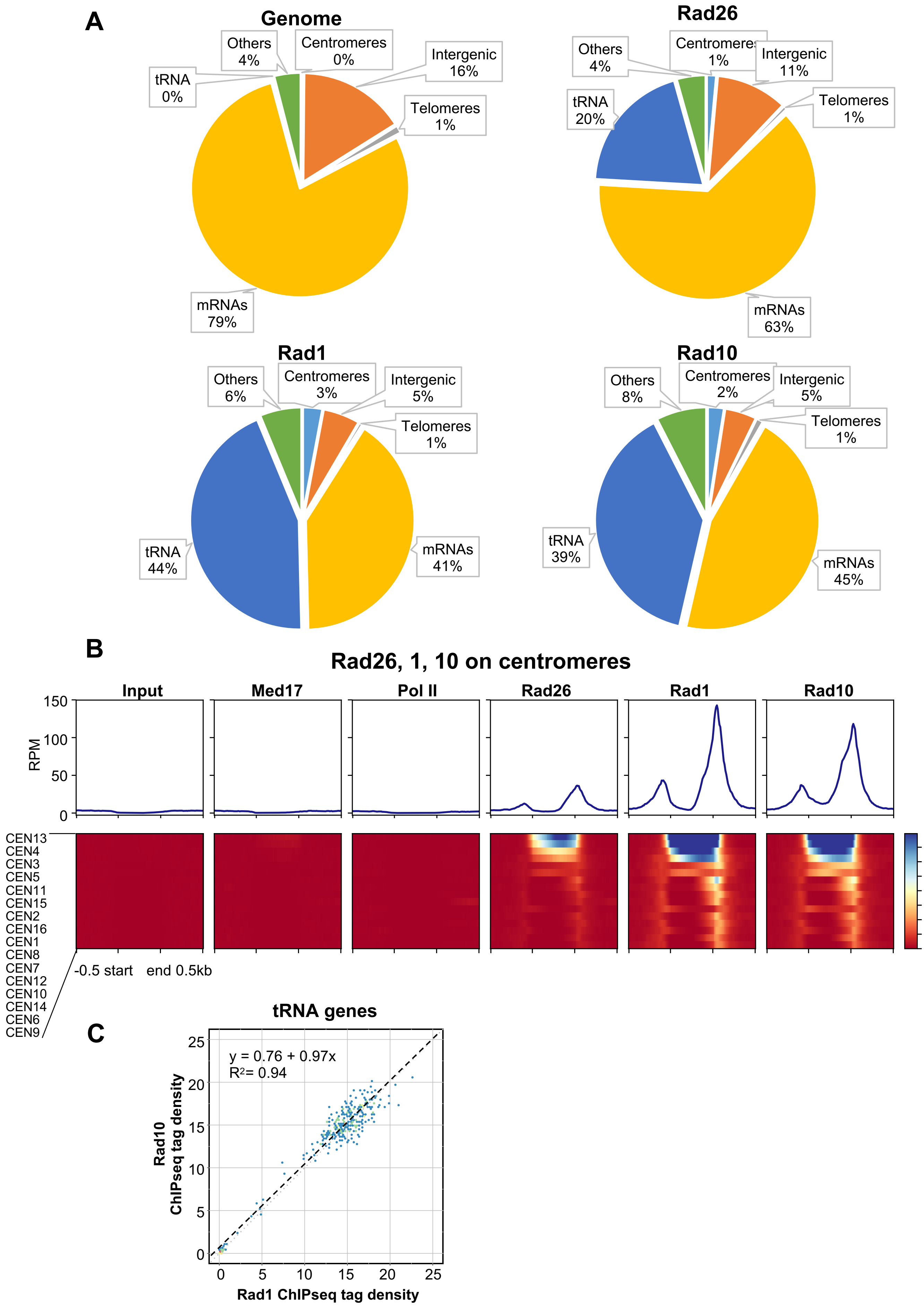

#### Supplementary Figure S4

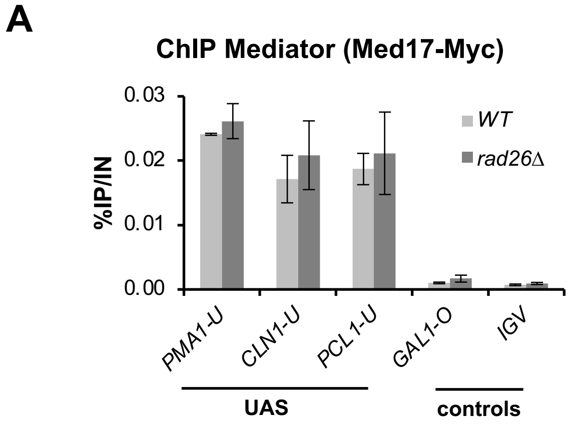


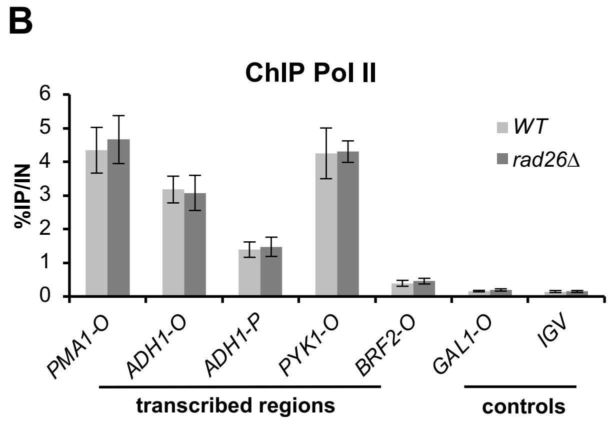


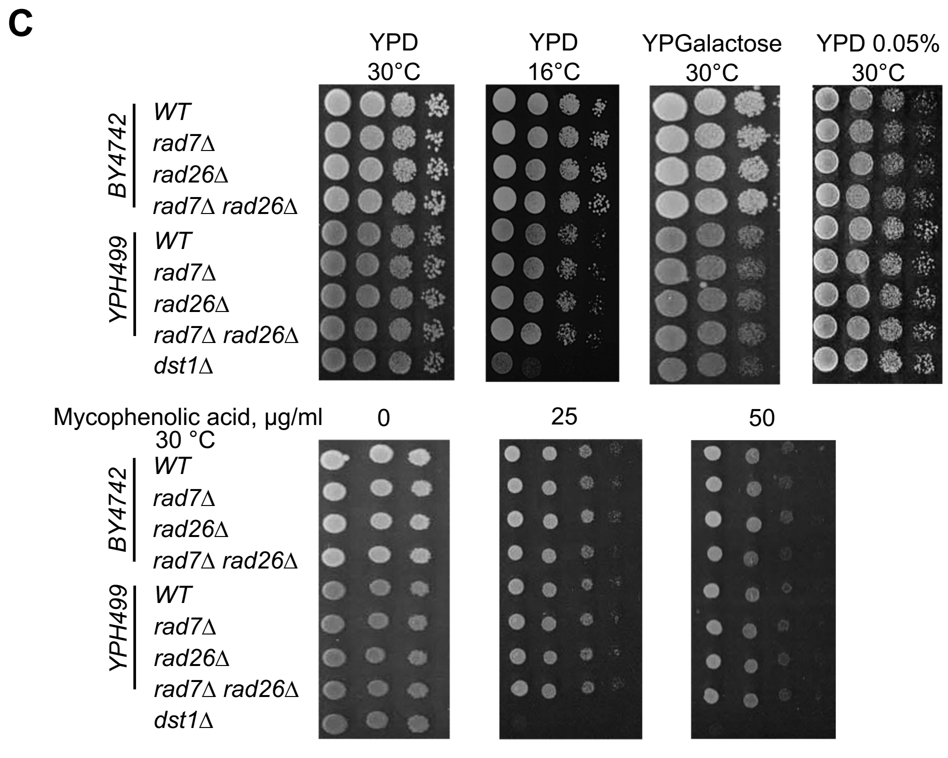

#### Supplementary Figure S5

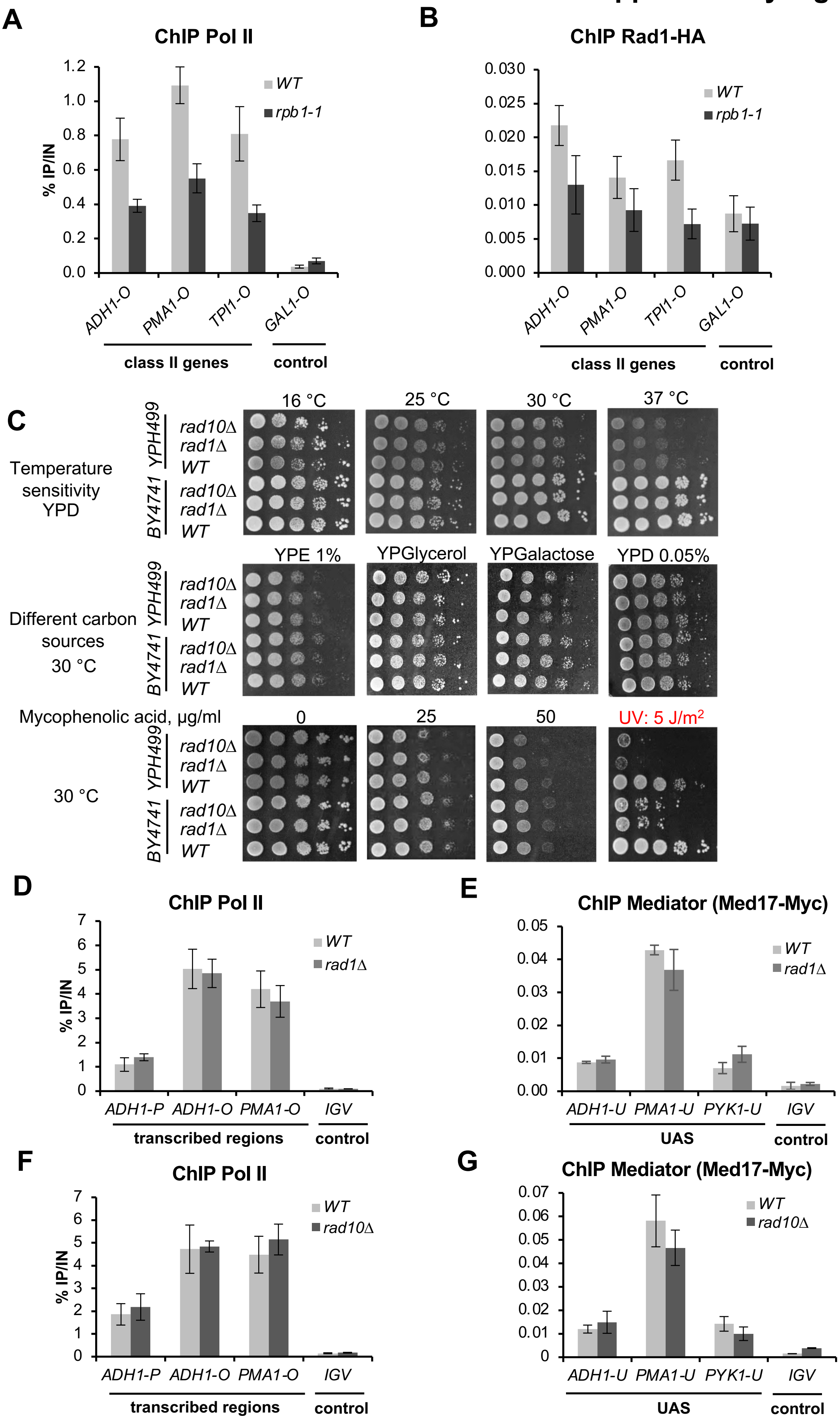

#### Supplementary Figure S6

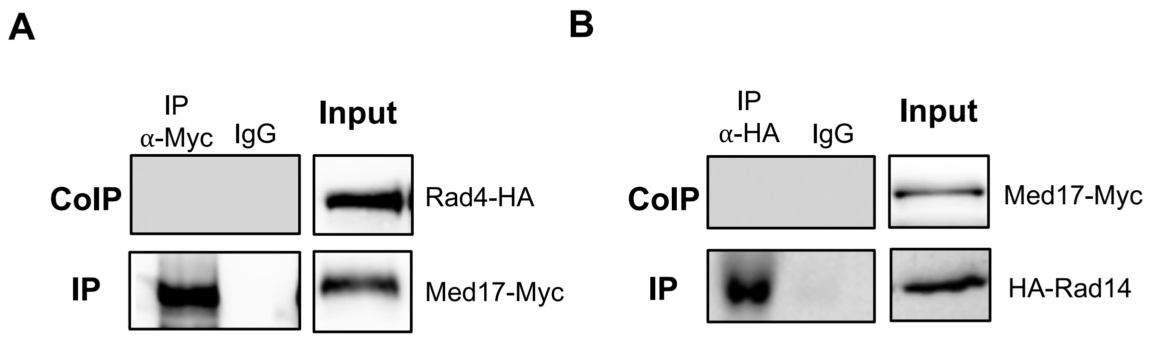

### Supplementary Figure S7

**A**

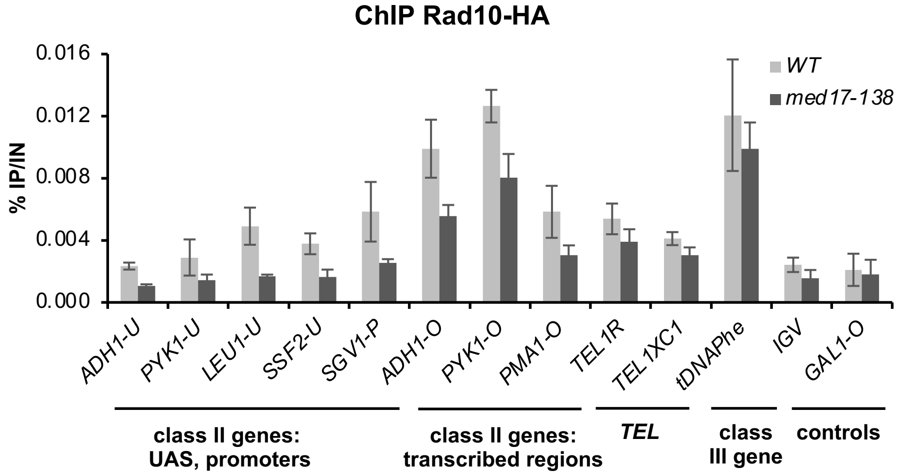


**B**

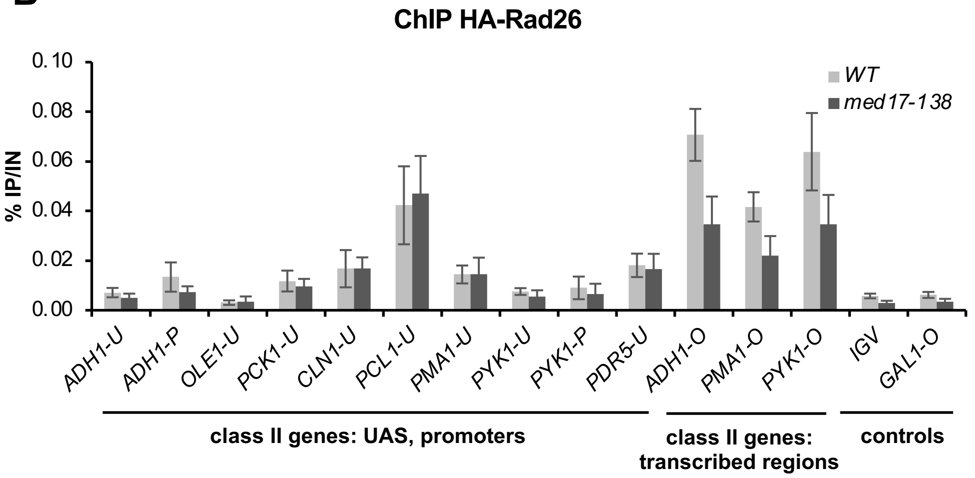

#### Supplementary Figure S8

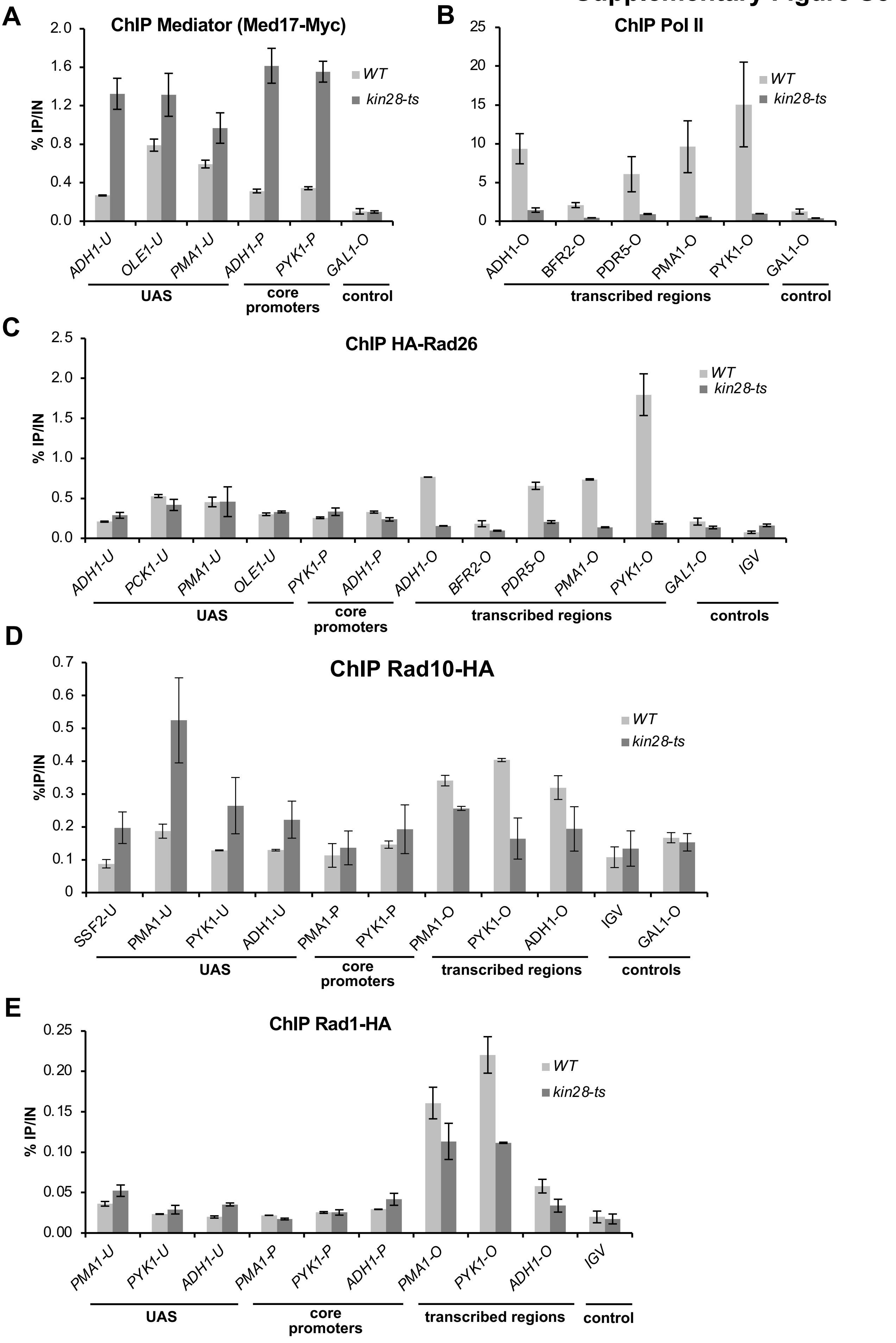

### Supplementary Figure S9

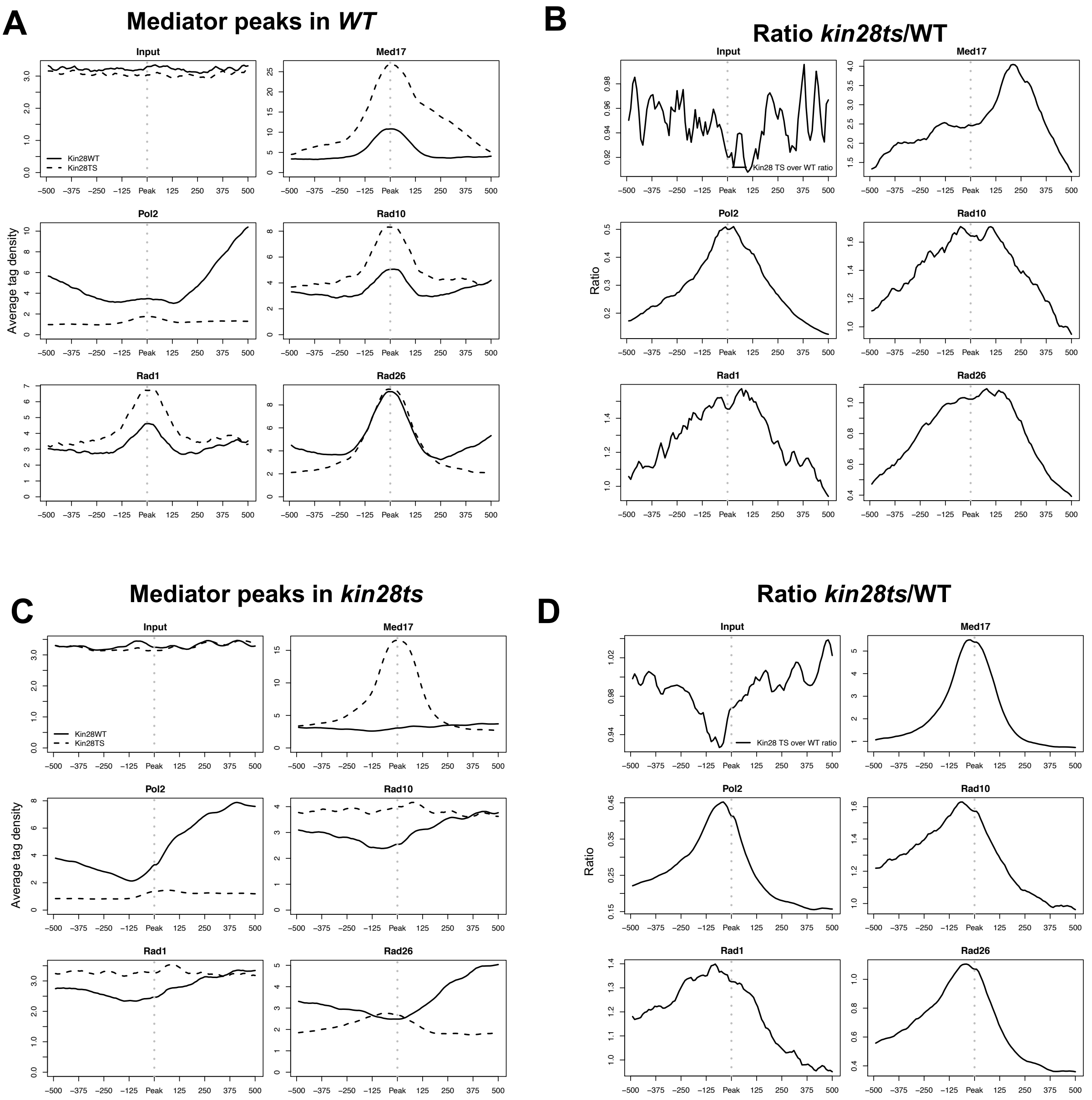

### Supplementary Figure S10

**A**

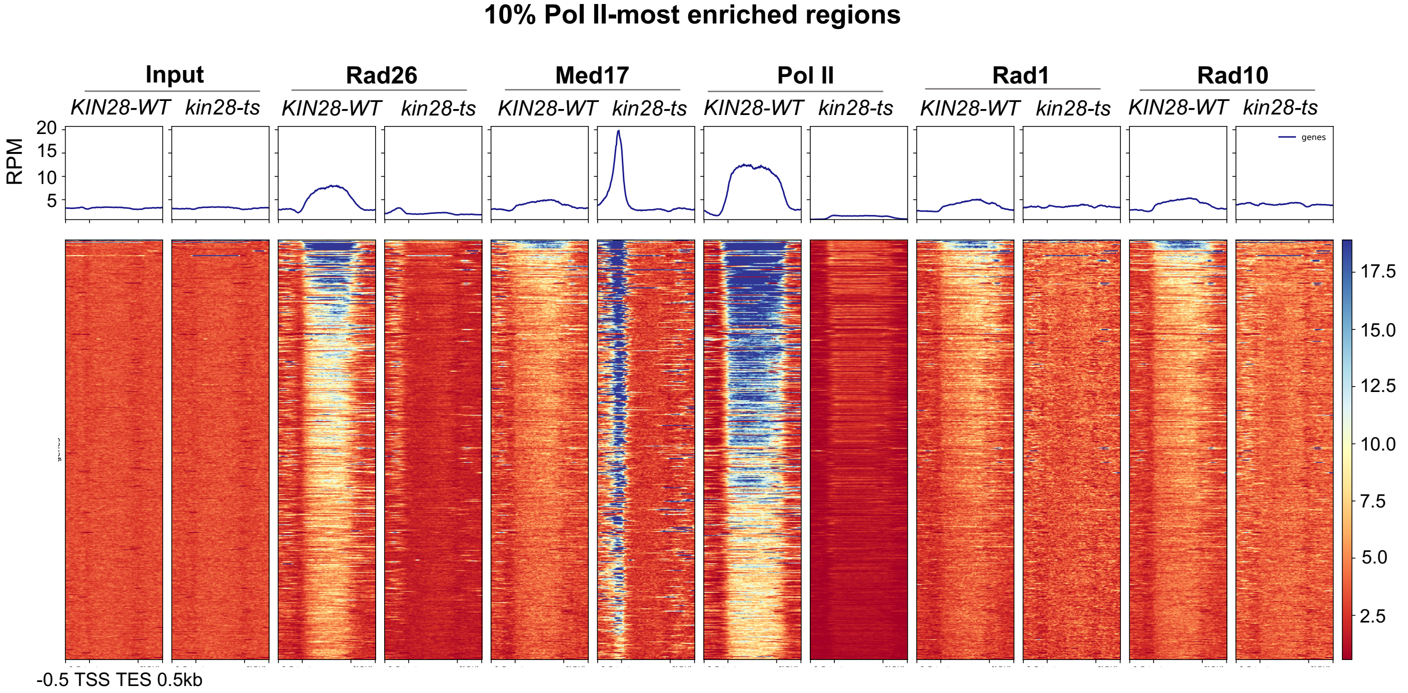


**B**

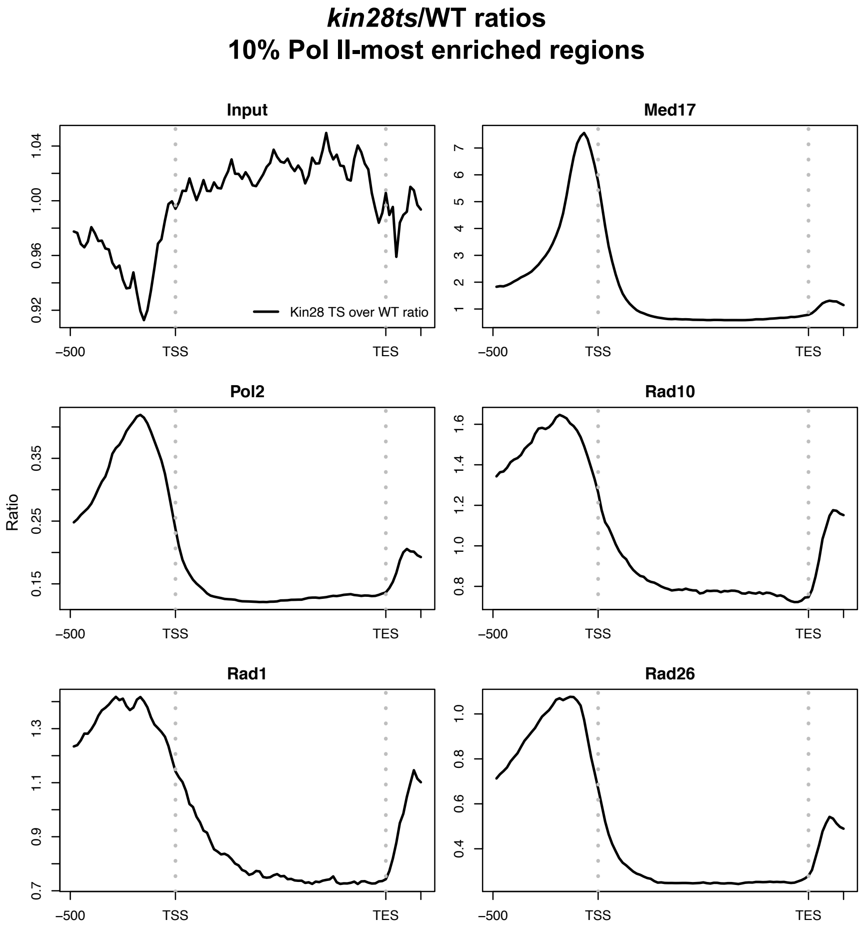

#### Supplementary Figure S11

**A**

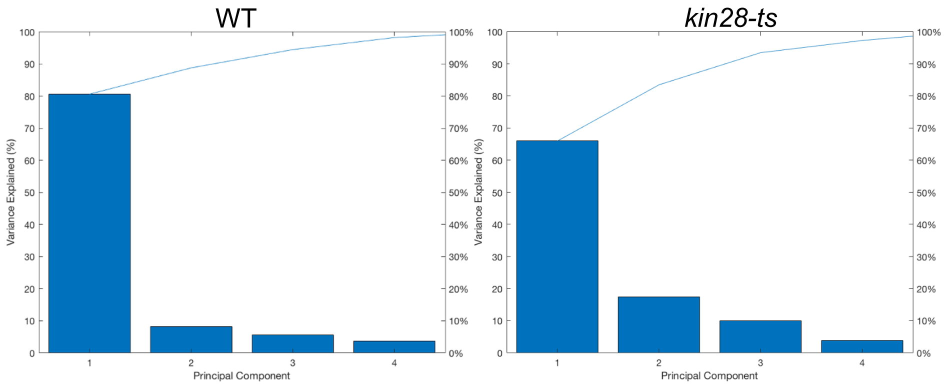


**B**

| | Norms WT | Norms *kin28-ts* |
|---|---|---|
| Med17 | 0.847 | 0.850 |
| Pol II | 0.829 | 0.938 |
| Rad10 | 0.602 | 0.613 |
| Rad1 | 0.737 | 0.714 |
| Rad26 | 0.830 | 0.715 |

**C**

| | PC1 WT | PC1 *kin28-ts* | PC2 WT | PC2 *kin28-ts* | PC3 WT | PC3 *kin28-ts* |
|---|---|---|---|---|---|---|
| Med17 | 0.502 | 0.464 | **-0.858** | **0.745** | 0.108 | 0.479 |
| Pol II | 0.517 | 0.426 | **0.820** | 0.517 | 0.246 | **-0.742** |
| Rad10 | **0.785** | **0.802** | -0.029 | -0.597 | **-0.619** | 0.001 |
| Rad1 | **0.626** | **0.640** | 0.126 | **-0.654** | **-0.770** | -0.403 |
| Rad26 | **0.537** | **0.676** | -0.044 | -0.146 | **0.842** | **0.722** |

**D**

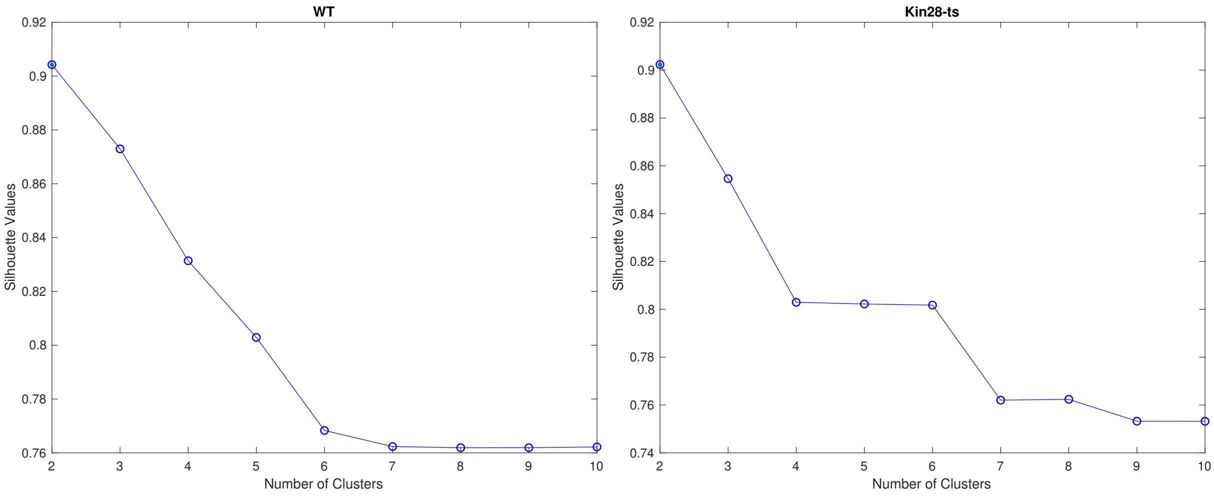

### Supplementary Figure S12

**A** ***WT, intergenic regions***

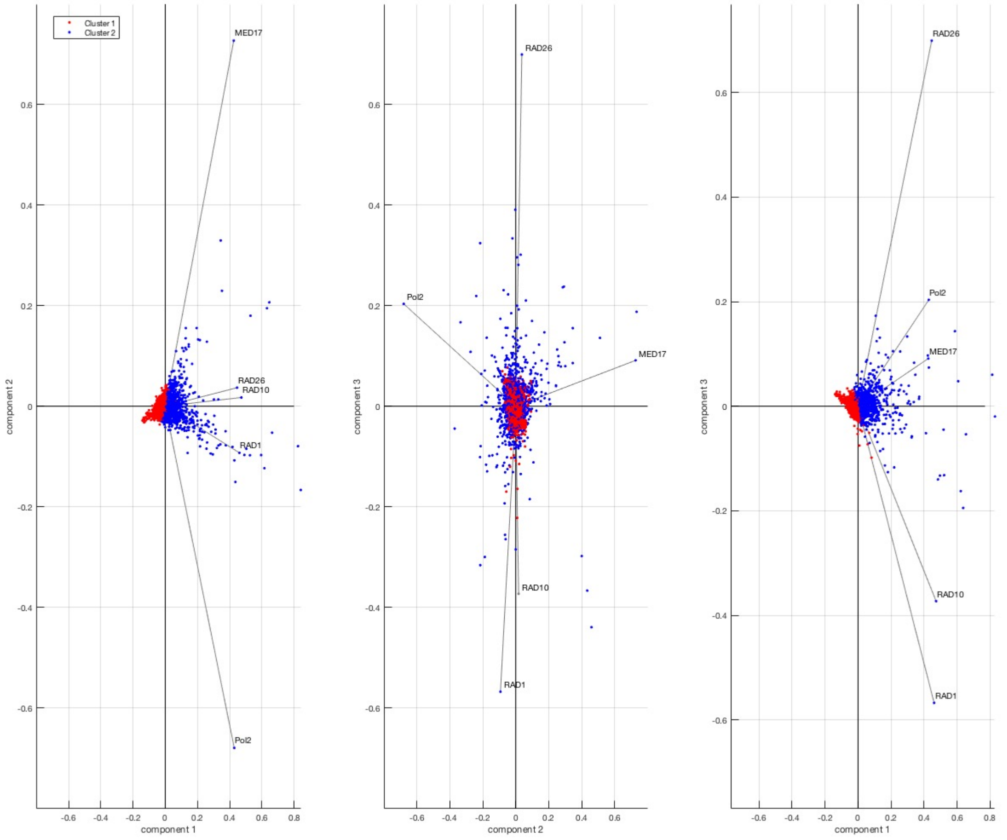


**B** ***kin28-ts, intergenic regions***

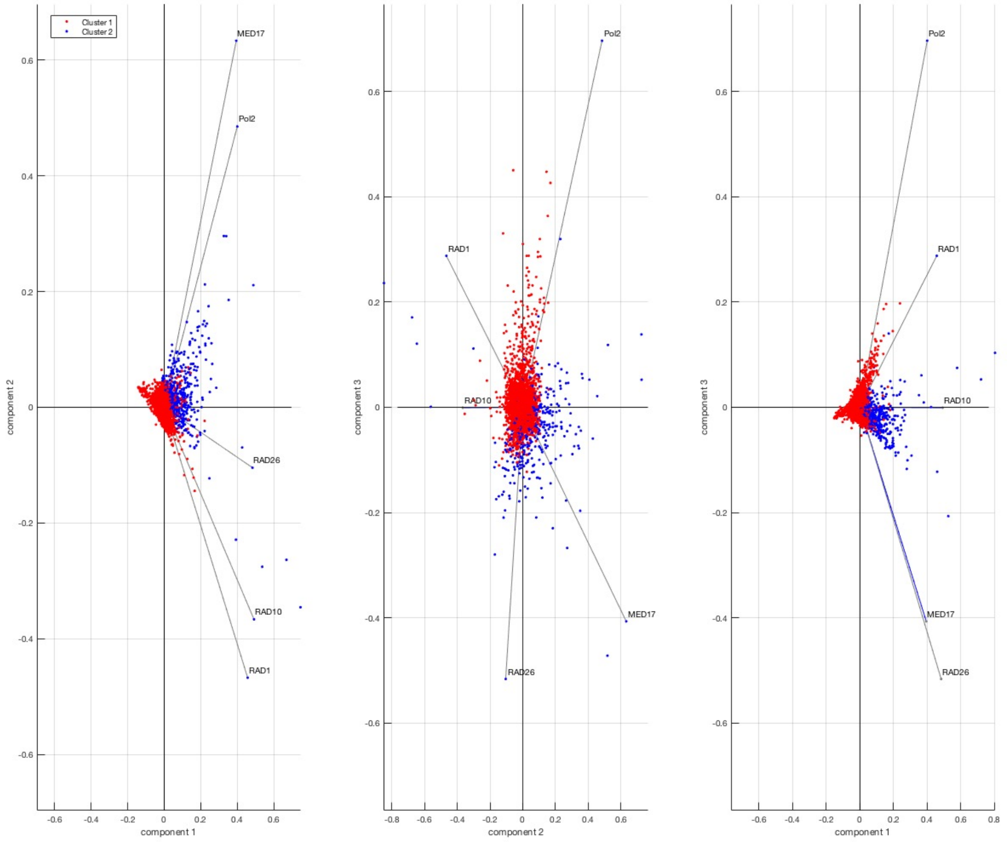

#### Supplementary Figure S13

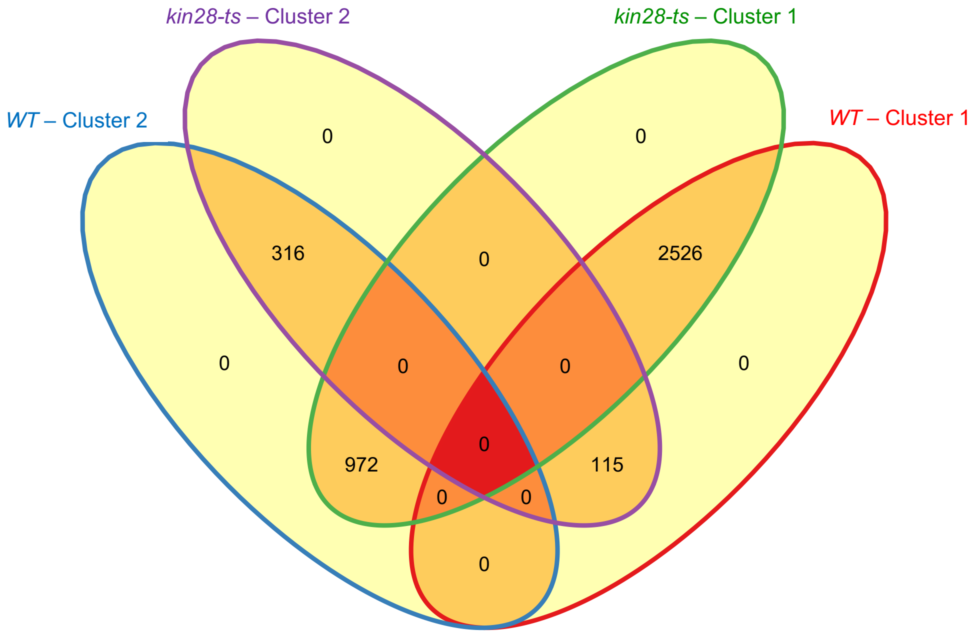
